# Autoimmune-like CD8⁺ T Cell Responses Drive Atherosclerotic Plaque Instability and Predict Cardiovascular Events

**DOI:** 10.64898/2026.04.26.720043

**Authors:** Jonathan Noonan, Nicholas Borcherding, Sander W. van der Laan, Nicholas A. Gherardin, Moustafa I Morsy, Joshua I. Gray, Lisa M. Domke, Anna M.D. Watson, Angela Huang, Anastasia Barbaro-Wahl, Shania A. Prijaya, Marcel Michla, Zahra Elahi, Yilu Huang, Nalin H. Dayawansa, Viktoria Bongcaron, Aidan P.G. Walsh, Prerna Sharma, Ana Maluenda, Peter Kanellakis, Gabriella E. Farrugia, Man-Kit Sam Lee, Andrew J. Murphy, Ian K. Hsu, Alexander R. Pinto, Chad J. Johnson, Yung-Chih Chen, Eleanor Eddy, Mélanie Le Page, Thomas M. Lovelock, Judy Wang, Michael Bourke, Zafreen Rahman, Vincent Varley, Joseph Kilby, Lorena Zentilin, Mauro Giacca, James D. McFadyen, Michal M. Mokry, Gerard Pasterkamp, Xiaowei Wang, Luciano Martelotto, Danika L. Hill, Thomas D. Otto, Robert A. Benson, Alex Bobik, Pasquale Maffia, Megan K.L. MacLeod, Thodur Vasudevan, Axel Kallies, Karlheinz Peter

## Abstract

**Background and aims:** Atherosclerotic plaque rupture is a major cause of myocardial infarction and stroke. However, the precise drivers of plaque destabilisation remain elusive. We hypothesised that antigen-driven, autoimmune-like T cell responses are central to the destabilisation and rupture of atherosclerotic plaques.

**Methods:** To dissect T cell responses specifically in unstable compared to stable plaques, we leveraged near-infrared autofluorescence (NIRAF) imaging–guided dissection of human carotid plaques. We also used our tandem stenosis model reflecting plaque instability as seen in patients to differentiate between unstable and stable plaques in mice. To explore T cell involvement, we studied T cell differentiation states and T cell receptor (TCR) repertoires by single-cell multi-omics. Then, testing if antigen-driven CD8^+^ T cell responses drive plaque instability in mice, we applied a combination of AAV8-PCSK9-induced atherosclerosis, tandem stenosis and TCR transgenic mice. Finally, we leveraged data from the AtheroExpress Biobank Study to link T cell immunity to histology-defined instability and cardiovascular outcomes.

**Results:** T cell responses in unstable versus stable atherosclerosis were distinct. Unstable human plaques contained highly expanded, autoimmune-like CD8⁺ T cells with markedly increased cytotoxic signatures, reduced exhaustion and distinct clonal repertoires compared to stable regions. Most plaque CD8⁺ T cells exhibited a pronounced tissue-resident transcriptional program. Moreover, the transcriptional signature of these plaque resident T cells was distinct from multiple other human tissues. Autoimmune-like cytotoxic and tissue-resident CD8^+^ T cell responses were also evident in murine atherosclerosis, where restricting the activation of antigen-driven CD8^+^ T cells prevented plaque destabilisation. Importantly, analysis of carotid endarterectomy samples from >1000 patients identified that intraplaque cytotoxic CD8⁺ T cell gene signatures strongly correlated with histological instability and predicted future strokes.

**Conclusions:** Integrated human, murine and clinical analyses demonstrate that autoimmune-like, cytotoxic CD8⁺ T cell responses are central drivers of plaque instability and major cardiovascular events. Targeting pathogenic CD8⁺ T cell responses may thus offer a compelling immunomodulatory strategy to stabilise plaques and reduce the risks of stroke and myocardial infarction.

**Graphical Abstract:** *Key Question:* Rupture of unstable atherosclerotic plaques is a typical cause of myocardial infarction and stroke. To understand the underlying cause and to prevent plaque rupture, we addressed the central hypothesis that autoimmune-like T cell responses drive plaque destabilisation and rupture.

*Key Findings:* CD8^+^ T cells are clonally expanded with increased cytotoxic signatures in unstable versus stable plaques (mice and humans) and require antigen recognition to drive plaque instability. Cytotoxic CD8^+^ T cell signatures in excised plaques correlate with increased future cardiovascular events.

*Take Home Message:* Autoimmune-like adaptive immune reactions, dominated by CD8^+^ T cells, are a major driver of plaque instability/rupture. Therefore, targeting pathogenic CD8⁺ T cell responses offers a compelling immunomodulatory strategy to stabilise plaques and reduce the risk of myocardial infarction and stroke. 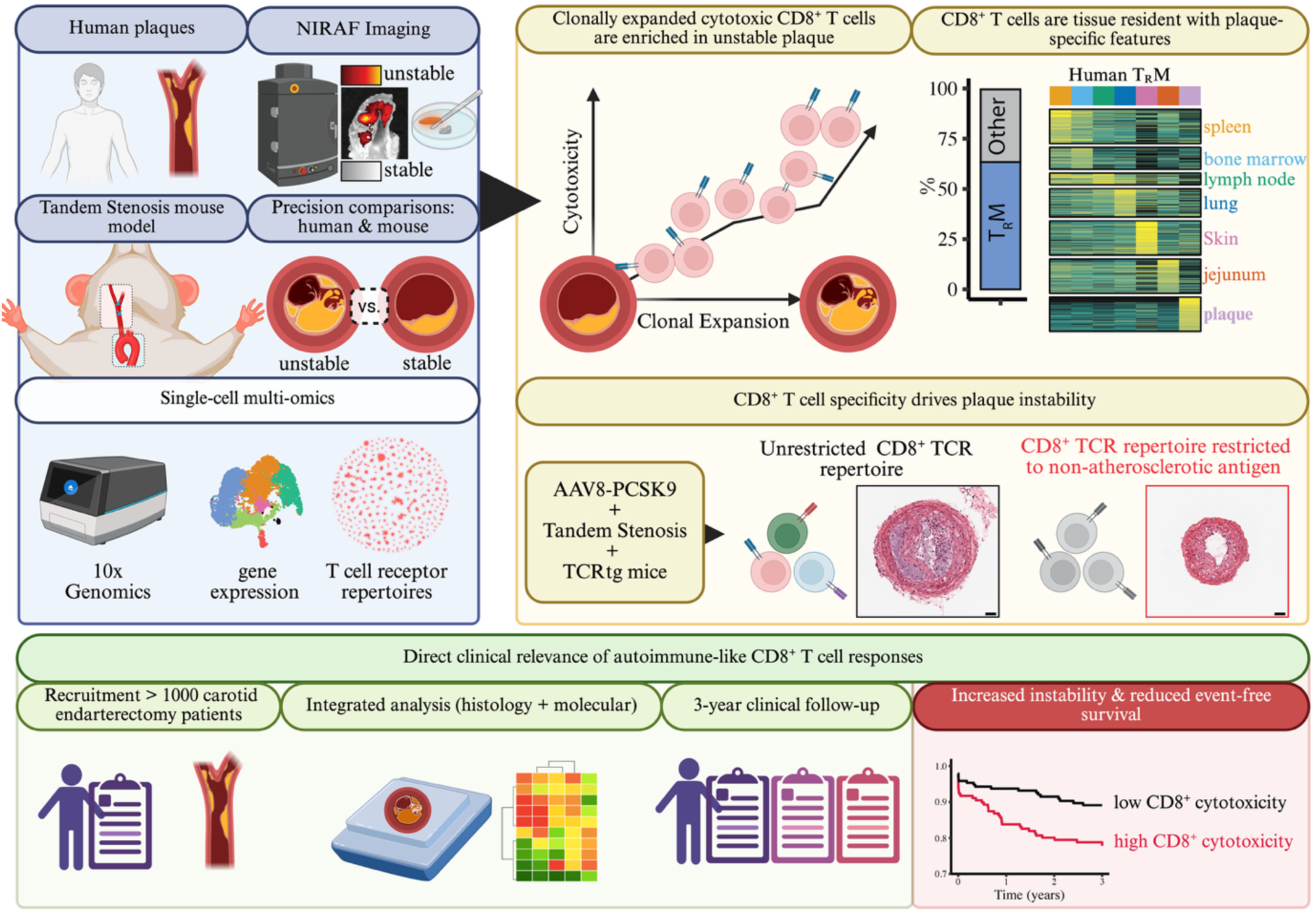

## Introduction

Atherosclerotic plaque rupture is the main cause of myocardial infarction and a major contributor to stroke, both leading causes of global morbidity and mortality.^1,2^ While modifying traditional cardiovascular risk factors, including hypercholesterolemia and hypertension, has significantly reduced event rates, many patients continue to suffer the catastrophic consequences of plaque rupture despite optimal therapy. Landmark clinical trials with anti-inflammatory drugs have now proven that inflammation is a major additional driver of cardiovascular risk.^3–6^ However, the overall application of anti-inflammatory therapeutics to prevent plaque rupture remains very limited, in part due to intolerable side effects including serious infection.^3–6^ Thus, there remains an inability to safely and explicitly address atherosclerotic inflammation to prevent plaque destabilisation.

T cells are the most abundant immune cell population in human atherosclerotic plaques.^7,8^ Several studies in mice and humans describe that T cells within plaques are heterogenous and typically possess pro-inflammatory phenotypes.^7,9–12^ Moreover, while not yet conclusive, emerging data suggest that T cells with predicted reactivity towards atherosclerosis-associated self-antigens clonally expand within plaques.^13–19^ This aligns with the hypothesis that atherosclerosis involves autoimmune or autoimmune-like mechanisms.^20^

Autoimmune(-like) responses are most comprehensively described in the context of apolipoprotein B (ApoB)-specific CD4^+^ T cells, which are present in humans and can drive plaque development in mice.^15–17,21–23^ This discovery of *de facto* self-reactivity builds on decades of evidence that pro-inflammatory CD4^+^ T cell function can drive plaque development.^24–30^ In contrast, CD4^+^ regulatory T cells (T_regs_) are known to robustly protect from atherosclerosis, highlighting a delicate balance between pro-atherogenic and protective T cell immunity.^24,31,32^ This understanding has recently been explored clinically, with the expansion of T_regs_ in patients using low-dose IL-2 demonstrated to reduce atherosclerotic inflammation in the IVORY trial.^33^ This work represents a major milestone for the cardiovascular field and T cell immunology, and highlights the therapeutic potential of modulating adaptive immunity to address atherosclerotic inflammation and risk.

While the mechanistic role of CD4^+^ T cells in plaque development is relatively clear, the role of CD8^+^ T cells is not.^24,34^ To date, both protective^35–37^ and pathogenic^19,38–40^ roles for CD8^+^ T cells have been described in plaque development, including via autoimmune-like mechanisms.^14,37^ Overall, existing data suggest that CD8^+^ T cell responses directed against cells integral to plaque stability (e.g., contractile smooth muscle cells) are likely detrimental,^40,41^ while those targeting pathogenic cells (e.g., activated smooth muscle cells, inflammatory macrophages and other antigen-presenting cells) may be protective.^35,36,42^ However, this question awaits more conclusive investigation.

Here, we explore the role of T cells in atherosclerotic plaque instability, ultimately focusing on the CD8^+^ compartment, and test if autoimmune-like mechanisms contribute to atherosclerotic disease in humans. We combine multi-omics approaches with analyses of precisely defined unstable versus stable human and murine atherosclerotic plaques, mechanistic preclinical modelling, and interrogation of a large, unique clinical cohort study with matched molecular, histological and outcome data.

## Results

### Human atherosclerotic plaque instability is associated with locally increased CD8^+^ T cell cytotoxicity

To define T cell responses in unstable compared to stable human atherosclerosis, we collected carotid plaques and matched blood from consecutively recruited patients undergoing carotid endarterectomy. Each plaque was then analysed using near-infrared autofluorescence (NIRAF) imaging, a unique technology that enables intrapatient identification, separation and then direct analysis of unstable (NIRAF^High^) versus stable (NIRAF^Low^) plaque. T cells were then independently isolated from unstable versus stable plaque regions and subjected to 10x genomics 5’ single-cell transcriptomics with paired αβ-T cell receptor (TCR) sequencing (scRNA-TCR-seq; **Figure 1A**). T cells from matched blood samples were also analysed to enable comparison between local (plaque) and peripheral (blood) T cells.

**Figure 1:**
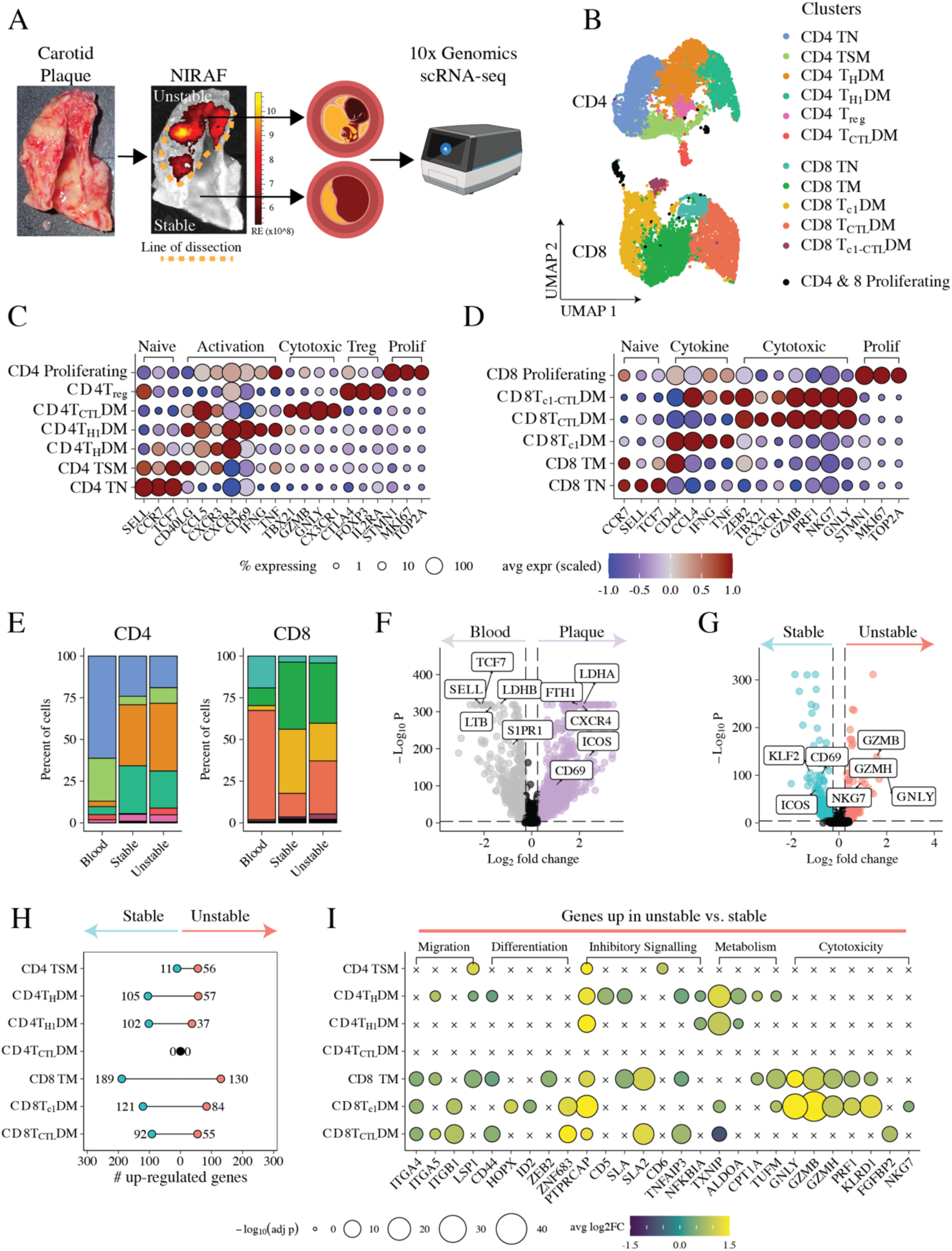
Distinct T cell signatures and increased cytotoxic profiles in human unstable versus stable atherosclerotic plaques. (A) Carotid plaques were isolated from patients undergoing endarterectomy and, using NIRAF imaging, were classified into unstable (NIRAF^High^) and stable (NIRAF^Low^) plaque segments. T cells were flow cytometrically enriched from unstable and stable segments and then subjected to 10x genomics scRNA-sequencing. Matched peripheral blood T cells were also included. (B) Annotated UMAP dimensionality reductions in CD4^+^ and CD8^+^ T cell transcriptomes. Select genes informing annotation of (C) CD4^+^ and (D) CD8^+^ T cells. (E) Proportions of CD4^+^ and CD8^+^ T cell clusters from blood, stable and unstable plaques. (F) Genes differentially expressed in plaque-derived versus blood-derived T cells. (G) Genes differentially expressed in unstable versus stable plaque-derived T cells. (H) Bubble plot presenting the number of genes differentially expressed by each cluster in unstable versus stable plaque-derived T cells and (I) the upregulation of select genes from this analysis. *n = 6* patients.

CD4^+^ and CD8^+^ T cells were computationally isolated using scGate^43^, clustered, and then annotated based on canonical gene expression in line with recently published guidelines with updated naming conventions (**Figure 1B**).^44^ In the CD4^+^ compartment, we identified clusters containing cells with naïve (TN; high expression of *CCR7, SELL* & *TCF7)* and memory phenotypes with expected access to secondary lymphoid organs (TSM; *SELL*, *CCR7, TCF7*, *CD40LG, CXCR3*). Two clusters contained disseminated helper memory cells (T_H_DM; *CXCR4*, *CCL5* with low/absent *SELL/CCR7*), one of which had a strong T_H_1 signature (T_H1_DM; *TNF, IFNG*). We also identified clusters containing cytotoxic (T_CTL_DM; *GZMB, GNLY*), regulatory (T_reg_; *FOXP3*, *IL2RA*) and proliferating (*MKI69*, *TOP2A*) CD4^+^ T cells (**Figure 1C & Supplementary Figure 1A**). In the CD8^+^ compartment, we identified clusters containing TN, T_CTL_DM and proliferating cells with gene profiles similar to CD4^+^ cells. In addition, we identified one memory cluster (TM; *CD44, NR4A3, GZMK*) where some cells may have access to secondary lymphoid tissues (via *CCR7* expression), one containing disseminated memory cells with evidence of type-1 cytokine production (T_c1_DM; *IFNG*, *TNF*) and one with a combined type-1 cytokine and cytotoxic signature (T_c1-CTL_DM; **Figure 1D & Supplementary Figure 1B**). Differentially expressed genes for each CD4^+^ and CD8^+^ cluster can be found in **Supplementary Tables 1 & 2**.

CD4^+^ and CD8^+^ clusters with signatures of effector function (e.g., cytokine production) were typically increased in plaque- versus blood-derived T cells (**Figure 1E**). This aligned with global (e.g., upregulated *CD69, ICOS, CXCR4*; **Figure 1F**) and cluster-specific (**Supplementary Figure 1C & D**) evidence of increased activation in T cells from plaques versus blood. Critically, we identified that CD8^+^ T_CTL_DM cells were enriched in unstable versus stable plaques, indicating a direct association between CD8^+^ T cells and plaque instability (**Figure 1E**). Moreover, cytotoxicity-associated genes were significantly upregulated in T cells from unstable versus stable plaques (*GZMB, GZMH, GNLY*; **Figure 1G, Supplementary Figure 1E**, **Supplementary Table 3**), with cluster-specific differential expression (**Figure 1H**) identifying this was driven by CD8^+^ T cells (**Figure 1I**). Thus, overall, we defined a transcriptional blueprint of CD4^+^ and CD8^+^ T cells in unstable compared to stable human plaques, which included a unique upregulation of cytotoxic CD8^+^ T cell profiles in unstable atherosclerotic plaques.

### Increased CD8^+^ T cell cytotoxicity and clonal expansion converge in unstable human plaques

We hypothesised that T cell responses promoting atherogenesis and plaque destabilisation are driven by antigen-specific and autoimmune-like mechanisms. Clonal expansion is a defining feature of such antigen-driven T cell responses.^45^ Previous data indicated clonal expansion of T cells in carotid plaques.^9,13,46^ However, these studies lacked paired αβTCR sequencing, which is necessary for accurate quantification of clonality, and/or did not include statistically robust analyses of multiple patients to enable generalisable conclusions and understanding of inter-individual variability. They also did not consider plaque instability. To overcome these limitations, we comprehensively assessed paired αβTCR repertoires and clonal expansion in human carotid atherosclerosis.

Clonally expanded T cells (detected ≥2 times within a patient) accounted for ∼25 to ∼65% of plaque T cells (**Figure 2A**). Chao indexes are commonly used to measure TCR repertoire diversity due to their resistance to sampling bias and sensitivity for rare clones, accounting for the extensive diversity of TCR repertoires versus the number of T cells it is possible to experimentally assess.^47^ Significantly reduced Chao indexes demonstrated reduced TCR diversity, indicating increased clonal expansion in atherosclerotic plaques versus blood consistent with an autoimmune(-like) response (**Figure 2B**). To infer whether plaque-infiltrated T cells respond to conserved or similar antigens across patients, we used Trex^48^ to cluster TCRs based on their sequence similarity and therefore their potential for shared specificity (**Figure 2C**). Plaque-derived T cells generated significantly larger TCR clusters than those from the blood, reflecting greater TCR repertoire similarity (**Figure 2C**). To validate this, we calculated Morisita indexes as an alternative measure of repertoire similarity.^47^ Increased repertoire similarity for TCRα (**Supplementary Figure 2A**) and β (**Supplementary Figure 2B**) chain usage in plaque versus blood further supported our conclusion that the same, or similar, antigens may drive plaque T cell responses across patients.

**Figure 2:**
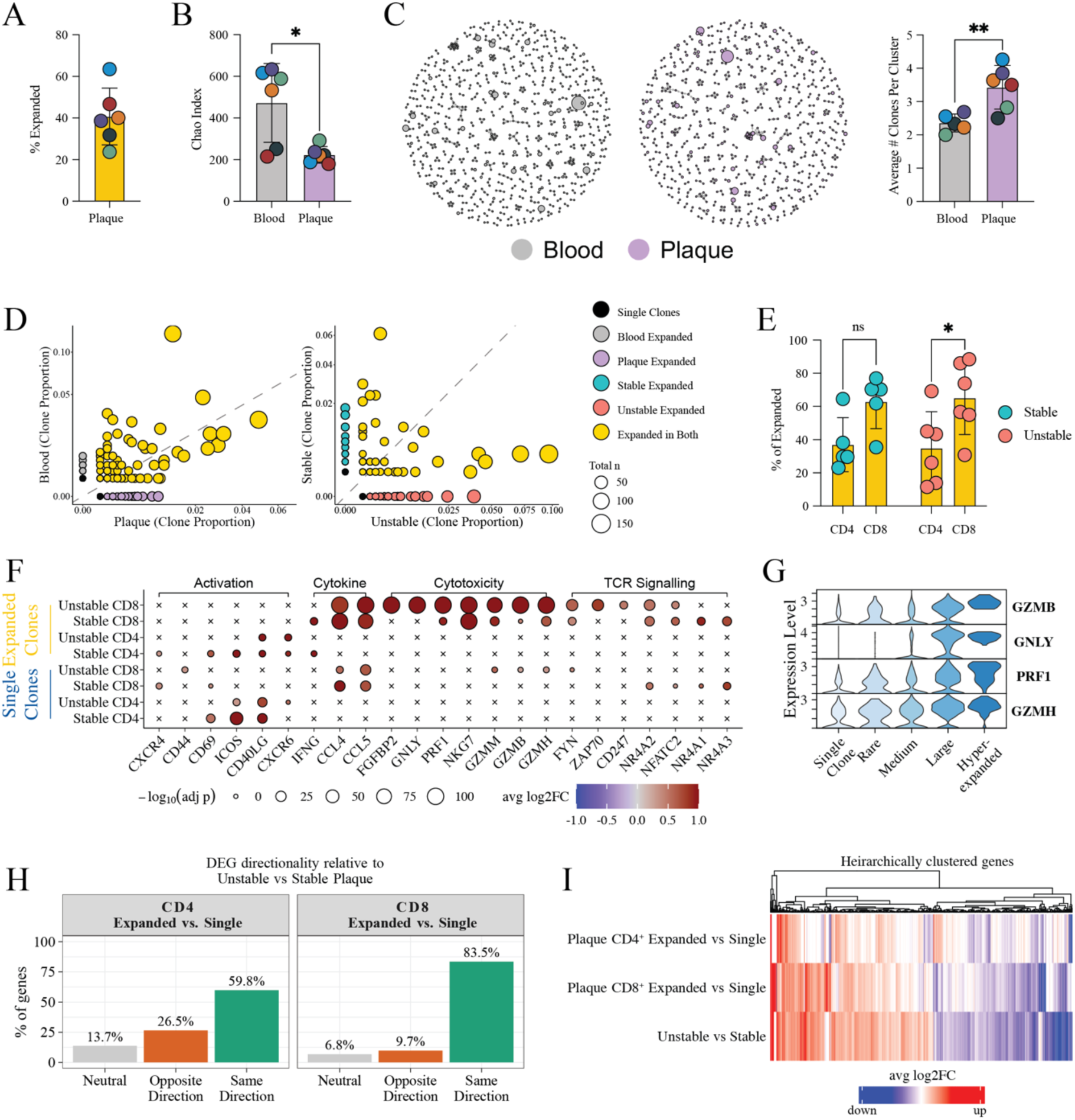
Clonal expansion of CD8^+^ T cells in human unstable versus stable atherosclerosis. T cells with both gene expression and T cell receptor (TCR) data from Figure 1 were isolated and the relationship between clonal expansion, T cell transcription and plaque instability assessed. (A) Percentage of T cells within human atherosclerotic plaques that were clonally expanded and (B) Chao index of plaque versus blood-derived T cells. (C) TCR amino acid edit distance-based clustering of blood (grey; left panel) and plaque (purple; right panel) T cells and assessment of the average number of clones per cluster. (D) Scatter plots identifying clones expanded in plaques versus blood (left panel) and unstable versus stable plaques (right panel). Plots identify clones unique to tissue origin, clones that were shared, and their overall abundance. (E) Percentage of CD4^+^ versus CD8^+^ T cells in stable versus unstable atherosclerotic plaque. (F) Select differentially expressed genes associated with T cell activation, cytotoxicity and TCR signalling from the comparison of expanded and single-clone CD4^+^ and CD8^+^ T cells from unstable versus stable plaques. (G) Violin plot of cytotoxicity-associated gene expression in CD8^+^ T cells stratified by cloneType. cloneTypes: Rare (≤ 0.01%), Medium (0.1% < X ≤ 1%), Large (1% < X ≤ 10%), Hyperexpanded (10% < X ≤ 100%). (H) Concordance of differentially expressed genes in unstable versus stable plaque T cells with those in CD4^+^ and CD8^+^ expanded versus single-clone T cells. Neutral genes did not change in either condition. Opposing genes showed opposite directionalities in unstable versus stable plaques and in CD4/CD8^+^ expanded versus single-clone analyses. Concordant genes exhibited the same directionality in unstable versus stable plaques and CD4/CD8^+^ expanded versus single-clone analyses. (I) Clustered heatmap showing concordance of genes upregulated in expanded versus single-clone CD4^+^ and CD8^+^ T cells and unstable versus stable plaque T cells. *n = 6* patients. Data were analysed using the Mann–Whitney test (B), Student’s t test (C), and ordinary two-way ANOVA (E). *\*p<0.05; **p<0.01*.

We next aimed to define differences in the clonal response between unstable versus stable atherosclerosis. First, the representation of expanded T cells varied across patients, without significant differences between unstable and stable plaques (**Supplementary Figure 2C**). Nevertheless, TCR repertoires were notably different in unstable versus stable plaques, highlighting T cell clones unique to each phenotype (**Figure 2D & Supplementary Figure 2D**). This suggests that autoimmune-like responses against different antigens are involved in unstable versus stable disease. We also found that CD8^+^ T cells were more expanded than their CD4^+^ counterparts in both unstable and stable plaques, although this was statistically significant only in unstable plaques (**Figure 2E**).

If T cells actively respond to antigens within plaques to promote instability, specific pro-inflammatory effector signatures in expanded T cells from unstable plaques would be expected as opposed to their single-clone and stable plaque counterparts. Consistent with this idea, we found that plaque instability and expansion significantly and concomitantly correlated with gene expression in both CD4^+^ and CD8^+^ T cells. Importantly, expanded CD8^+^ T cells were the major source of instability-related cytotoxic signatures (**Figure 2F**). Expanded CD8^+^ T cells from unstable plaques were also uniquely enriched for *ZAP70* and co-expressed several other genes involved in or downstream of TCR signalling (e.g., *FYN, CD247*, *NR4A2*), suggestive of ongoing or recent antigen recognition. Expression of cytotoxicity-associated genes in plaque-derived CD8^+^ T cells also increased with the magnitude of expansion (**Figure 2G**).

To further evaluate the relationship between clonal expansion and plaque instability, we compared gene expression in expanded versus single-clone CD4⁺ and CD8⁺ T cells. We then contrasted these genes to those differentially expressed between unstable and stable plaques. Genes upregulated in expanded CD8⁺ T cells showed greater concordance with the unstable plaque signature (83.5%) than those in CD4⁺ T cells (59.8%) (**Figure 2H & I**). Taken together, these data connect CD8⁺ T cell clonal expansion, antigen recognition and cytotoxicity in unstable plaques, supporting the conclusion that autoimmune(-like) CD8^+^ T cell responses are directly involved in plaque instability.

### Plaque instability is associated with reduced T cell exhaustion

T cell exhaustion represents a mechanism for preventing excessive T cell–mediated inflammation, with exhausted T cells (TX) characterised by distinct transcriptional/proteomic profiles and diminished effector function. In cancer, infection and autoimmunity, it is well established that chronic antigen recognition and persistent TCR signalling induce T cell exhaustion.^49^ Thus, exhaustion would be expected if atherosclerosis is mediated by chronic autoimmune mechanisms, but could serve to reduce inflammation and promote plaque stability. Although previous studies identified exhaustion-associated genes (e.g., *TOX*) and proteins (e.g., PD-1) in T cells within human plaques^9,13,50^, whether exhaustion predominantly affects CD4⁺ or CD8⁺ T cells and whether TX are enriched in plaques or associate with plaque (in)stability remain to be determined.

Consistent with strong ongoing antigen recognition,^51–53^ TCR signalling–associated genes were significantly increased in CD8^+^ versus CD4^+^ plaque T cells (**Supplementary Figure 3A**). *TOX,* encoding a transcription factor essential for exhaustion,^54–57^ was also predominantly restricted to CD8^+^ T cells in plaques (**Figure 3A**). These data indicate that exhaustion is restricted to CD8^+^ cells, which we focused on in subsequent analyses. To accurately identify CD8^+^ TX in our data, we used two complementary approaches. First, we defined T cells as exhausted if they expressed *TOX* alongside ≥3 additional exhaustion-associated genes (*PDCD1, LAG3, HAVCR2* [encodes TIM3], *TIGIT*, *IRF4* or *ICOS*). Second, we leveraged ProjectTILs^58^ to map CD8^+^ T cells in our data against a robust reference dataset containing prototypical CD8^+^ TX. Both approaches enabled the identification of CD8^+^ T cells with strong co-expression of exhaustion-associated genes (**Figure 3B & Supplementary Figure 3B**), with most TX present within the CD8^+^ TM and T_C1_DM clusters (**Figure 3C & D; Supplementary Figure 3C**). Consistent with exhaustion typically being localised to chronically inflamed tissues where persistent TCR stimulation occurs,^57^ we found CD8^+^ TX were near-absent in the blood but significantly increased in plaques (**Figure 3E**). We performed linear regression to confirm that the stereotypical relationship between TCR-signalling and exhaustion persists in plaque CD8^+^ T cells,^57,59^ validating that antigen recognition is a vital component of exhaustion in atherosclerosis (**Figure 3F, Supplementary Figure 3D**).

**Figure 3:**
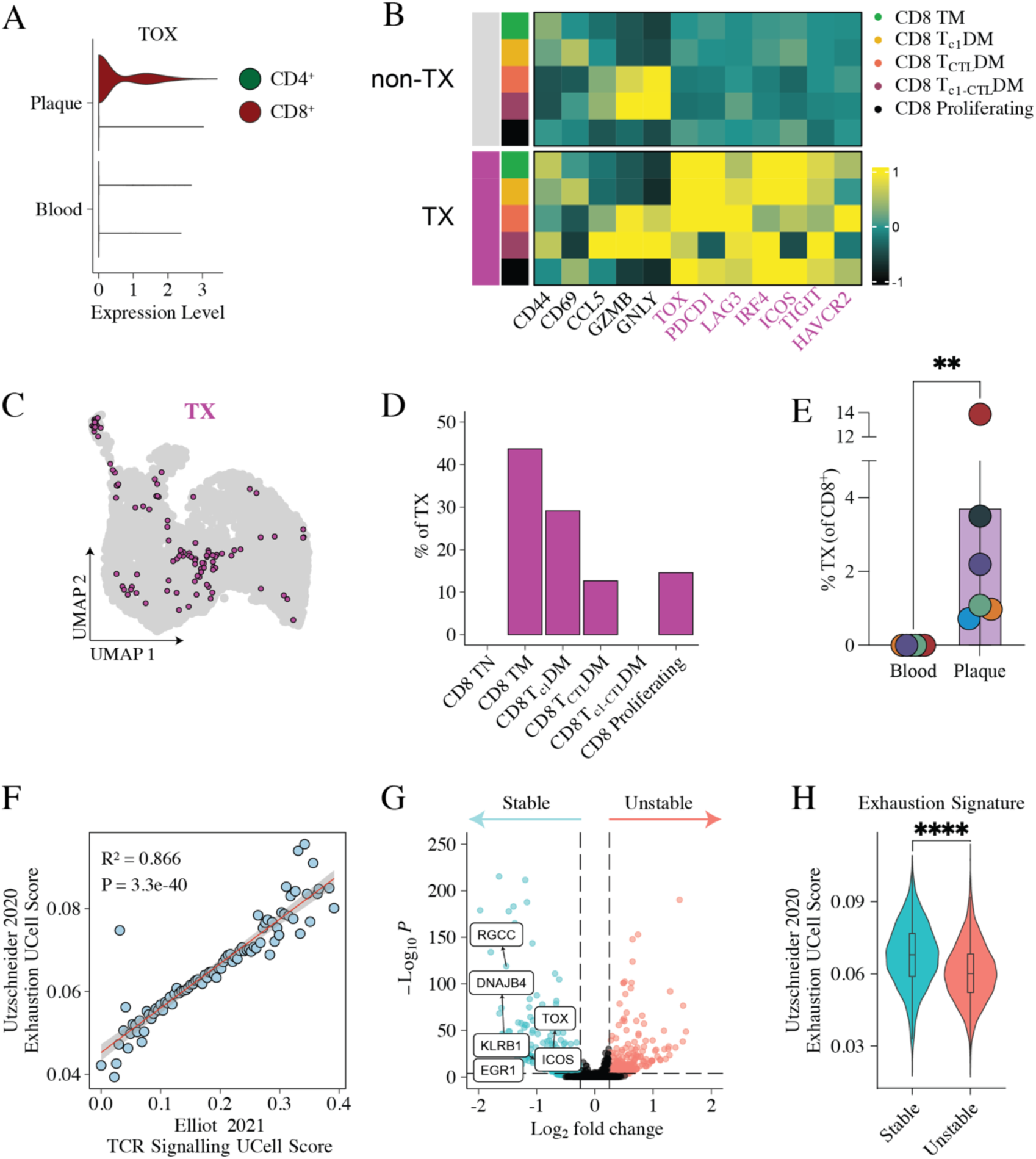
T cell exhaustion is CD8^+^ T cells increased in human plaques but reduced in unstable atherosclerosis. Data from Figure 1 were further analysed in the context of T cell exhaustion. (A) Violin plot of *TOX* expression in blood- and plaque-derived CD4^+^ versus CD8^+^ T cells. As *TOX* expression was confined to CD8 T cells, all further analyses focused only on the CD8 compartment. (B) Heatmap of select genes, including exhaustion-associated genes (highlighted in purple), in manually annotated TX vs. non-TX for each CD8^+^ T cell cluster. (C) UMAP positioning of TX identified using reference mapping and (D) the percentage of TX coming from each CD8^+^ T cell cluster. (E) Percentages of CD8^+^ TX cells in plaques versus blood based on reference mapping. (F) Correlation between T cell exhaustion gene-set scoring based on Utzschneider *et al*.,^63^ and TCR signalling based on Elliot *et. al*.,.^51^ in plaque CD8^+^ T cells. (G) Volcano plot of differentially expressed genes in unstable versus stable plaque CD8^+^ T cells, highlighting select genes associated with T cell exhaustion. (H) T cell exhaustion scoring in unstable versus stable plaque CD8^+^ T cells. *n* = 6 patients. Data were analysed using the Mann–Whitney test (E & H). *\*\*p<0.01; ****p<0.0001*.

We next asked whether exhaustion is altered in unstable versus stable disease. Comparing CD8^+^ T cells in unstable versus stable plaques, we noted that stable plaque CD8^+^ T cells were significantly enriched for exhaustion-associated genes including *TOX* (**Figure 3G**, **Supplementary Table 4**). CD8^+^ T cells from stable plaques also had significantly higher exhaustion gene-set scores than those from unstable plaques (**Figure 3H**). Together, these data indicate that CD8^+^ T cell effector function may be reduced in human stable versus unstable plaques due to greater T cell exhaustion. Thus, promoting exhaustion could provide a means of dampening T cell–mediated plaque inflammation, as has been suggested in the context of autoimmunity.^60–62^ Likewise, reduced exhaustion in unstable plaques aligns with increased CD8^+^ T cell effector function.

### Unstable and stable plaques contain tissue resident-like CD8+ T cells with plaque-specific transcriptional profiles

Our combined evidence supported that clonally expanded CD8^+^ T cells with increased cytotoxicity are a hallmark of unstable plaques. To expand on this and understand the transcription factor regulons underpinning plaque CD8^+^ T cell responses, we used SCENIC-based^64^ gene regulatory network (GRN) analysis. Notably, we detected increased activity of regulons implicated in T cell tissue residency^65–69^ relevant to both unstable and stable plaque versus blood-derived CD8^+^ T cells. These included increased activity of BHLHE40 and FOS accompanied by concomitant reductions in known negative regulators of tissue-resident memory T cell (T_R_M) differentiation (e.g., *EOMES*, *TBX21*, *KLF2*, *TCF7*; **Figure 5A** & **Supplementary Table 5**).

T_R_M provide long-lasting local protection within tissues but can also propagate chronic inflammation and autoimmunity.^70,71^ Current data indicate that T_R_M are rare within atherosclerotic plaques.^72,73^ However, our SCENIC analysis suggested that plaque CD8^+^ T cell residency may in fact be far more common. One reason for this discrepancy could be that a small number of genes or proteins implicated in residency in other tissues have been directly extrapolated to the atherosclerosis context (e.g., *ITGAE*, *ITGA1*, *ZNF683*).^9,72–74^ However, T_R_M typically possess tissue and context-specific features,^67,68,75–77^ which have yet to be identified in atherosclerosis. We therefore aimed to define the transcriptional identity of human plaque CD8^+^ T_R_M-like cells to enable their quantification and interrogation. Using gene-set scoring, we found that comprehensive T_R_M gene signatures from multiple independent studies, and the average of these, were all significantly increased in plaque versus blood CD8^+^ T cells (**Supplementary Figure 4A**). Projected onto our CD8^+^ T cell clusters, we found that CD8^+^ T_R_M signatures increased in all non-TN clusters in plaques versus blood but were strongest in plaque CD8^+^ TM and T_c1_DM (**Figure 4B**). Thus, these results validate a strong propensity towards CD8^+^ T_R_M programming in human plaques.

**Figure 4:**
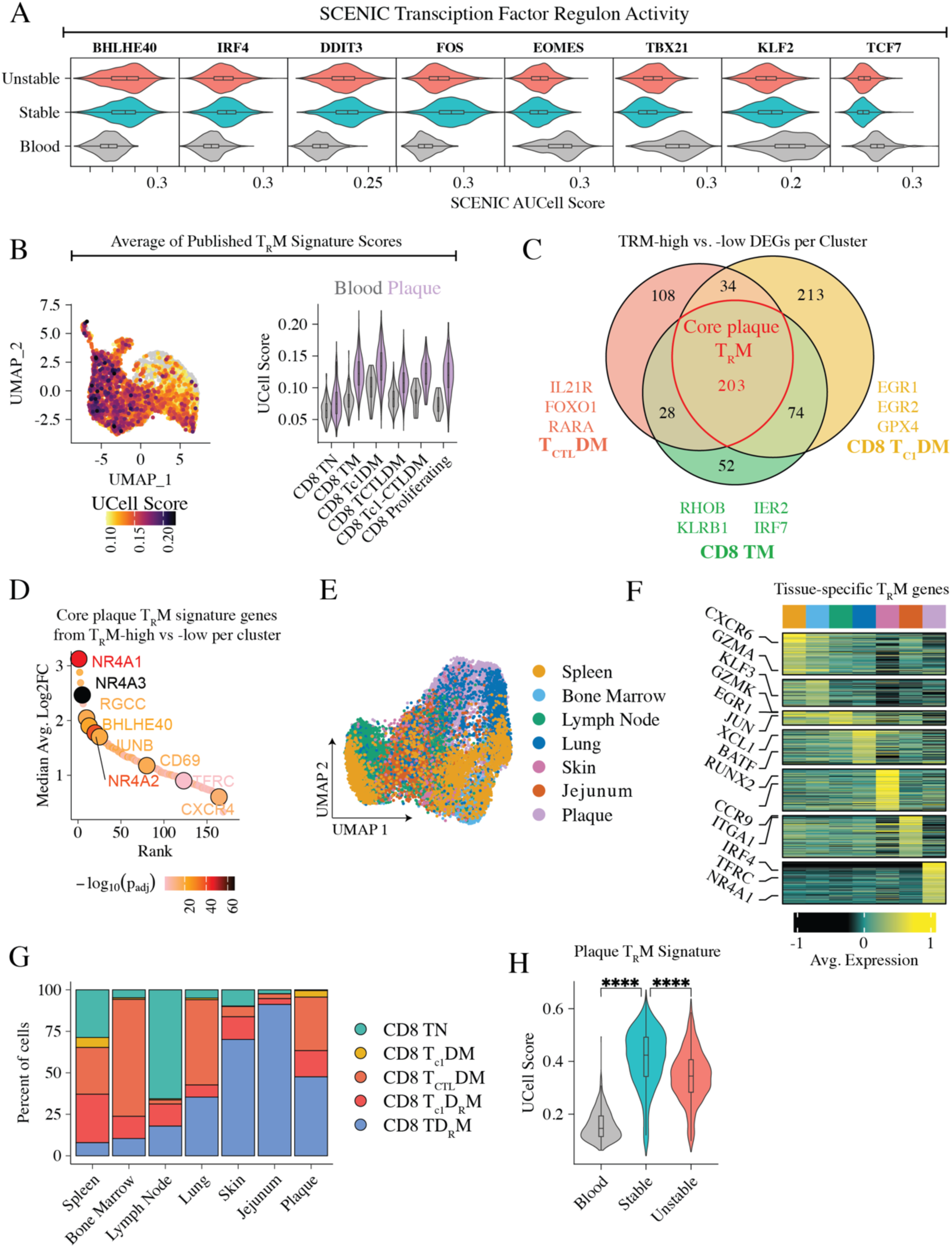
Human plaque CD8^+^ T cells are T_R_M-like with a plaque-specific transcriptional signature that is reduced in unstable versus stable disease. CD8^+^ T cell data from Figure 1 were further analysed in the context of tissue residency. We performed SCENIC analysis to infer the activity of GRNs (regulons) in T cells derived from human unstable plaques, stable plaques and blood. (A) Violin plot of select regulon activity associated with T cell tissue residency. (B) Feature and Violin Plot demonstrating the average UCell score from established tissue-residency gene sets from multiple published sources in relation to cluster and tissue origin. (C) CD8^+^ T cells identified as T_R_M-high versus T_R_M-low (upper versus lower quartiles of average T_RM_ score) were then compared and overlapping genes for the major effector clusters are presented. (D) Rank-plot of the 203 genes commonly upregulated in plaque T_R_M-high cells considered to be the core signature for human plaque T_R_M. Genes were ranked based on their median Avg. Log2FC and select genes are labelled. (E) Data from CD8^+^ plaque T cells were integrated with CD8^+^ T cell data isolated from multiple tissues in Poon *et. al.,.* UMAP dimensionality reduction of CD8^+^ T cells from integrated data highlighting tissue origin of each cell. (F) Select genes differentially upregulated by CD8^+^ T_R_M-high cells across tissues. Genes also upregulated by T_R_M-intermediate or low cells were excluded. (G) Clusters from the integrated dataset were annotated to identify tissue resident CD8^+^ T cell clusters. The representation of each subset across each tissue, including plaques, are shown. (H) Gene-set scoring of core plaque T_R_M genes in blood versus stable and unstable atherosclerotic plaques. *n = 6* patients. Data were analysed using an ordinary one-way ANOVA (H), *\*\*\*\*p<0.0001*.

To then identify which T_R_M signature genes are relevant to plaque T cells we compared CD8^+^ T cells with a high (upper quartile) versus low (lower quartile) average T_R_M score. Of T_R_M genes reported in at least two of the gene sets used, we found 128 of these (e.g., *NR4A1*, *JUNB*, *CD69 and BHLHE40*) were significantly upregulated in plaque T_R_M-high cells (**Supplementary Figure 4B**). Moreover, 70 genes (e.g., *ATF3, RGCC* and *CXCR4*) were upregulated in plaque T_R_M-high cells that were absent or inconsistently reported elsewhere, suggesting these may be of increased relevance specifically to plaque residency. To define a broadly applicable and refined plaque T_R_M signature while also identifying cluster-specific features of residency, we repeated the T_R_M-high versus T_R_M-low comparison within each cluster (**Figure 4C**). Genes previously used to identify T_R_M in human plaques (e.g., *ITGAE*, *ITGA1*, *ZNF683*) were notably not associated with T_R_M-high cells. However, plaque CD8^+^ T_R_M-high cells consistently upregulated 203 genes including *NR4A1-3, RGCC, BHLHE40, JUNB* and *CD69* versus T_R_M-low cells, regardless of their cluster of origin (**Figure 4D** & **Supplementary Table 6**).

To then identify transcriptional programs enriched in plaque CD8^+^ T_R_M versus other tissues, we integrated our plaque scRNA-seq data with similar analyses of other organs from Poon *et al.*,^75^ in which T_R_M are highly represented and well characterised (**Figure 4E**; including spleen, bone marrow, lymph node, lung, skin and jejunum). As expected, CD8^+^ T_R_M-high cells were transcriptionally heterogenous across tissues (**Figure 4F**). Importantly, plaque T_R_M-high cells possessed unique features, including the upregulation of *TFRC*, suggesting adaption to the iron-rich environment of plaques. Plaque T_R_M-high cells were also notably enriched for exhaustion-associated genes including *TOX*, *IRF4, NR4A1* and *ICOS* versus other tissues (**Figure 4F**, **Supplementary Figure 4C** & **Supplementary Table 7**). T_R_M populations were thought to form and stabilise predominantly after antigen has been cleared.^78–80^ However, recent studies have identified that persistent antigen stimulation leads to the differentiation of CD8^+^ T_R_M with exhausted phenotypes.^65,81,82^ Thus, our results now extend this observation to the cardiovascular disease setting by identifying exhausted T_R_M cells in atherosclerotic plaques.

To quantify plaque CD8^+^ T_R_M we next annotated our integrated multi-tissue dataset with specific consideration for tissue residency. In addition to TN, T_c1_DM and T_CTL_DM states described previously, we identified several clusters of disseminated T_R_M cells expressing canonical and plaque-related T_R_M-associated genes (**Supplementary Figure 4D** & **E**). This included clusters of T_R_M with type-1 cytokine (T_c1_D_R_M) and cytotoxic gene-expression (T_CTL_D_R_M). The representation of T_R_M in tissues from Poon *et. al*., in our analysis was highly consistent with the original study,^75^ with the jejunum and skin having the largest T_R_M compartments (**Figure 4G**). Critically, >50% of plaque CD8^+^ T cells possessed a tissue-resident transcriptional state. CD8^+^ T_R_M-high cells also displayed increased cytotoxicity in unstable versus stable plaques (**Supplementary Figure 4F**), indicating their involvement in destabilisation. However, we also noted that the overall CD8^+^ T_R_M signature was reduced in the unstable versus stable plaque, suggesting increased prevalence of newly recruited and/or non-resident cells (**Figure 4H**). Overall, these data reveal that CD8 T_R_M with plaque-specific transcriptional features are a major population in atherosclerotic plaques, that CD8^+^ T_R_M have an increased cytotoxic signature in unstable versus stable plaques, and that additional CD8^+^ T cell recruitment may play an important role in driving plaque instability.

### Unstable plaques in tandem stenosis mice contain clonally expanded, cytotoxic, tissue resident and exhausted T cells

To enable preclinical mechanistic testing, we employed our previously developed tandem stenosis (TS) mouse model, which reliably recapitulates key features of the unstable atherosclerotic plaques observed in patients with myocardial infarction.^83,84^ T cells were substantially more activated in unstable compared to stable plaques in the TS model (**Supplementary Figure 5A**). T cells were sorted from TS unstable plaques and subjected to scRNA-TCR-seq analysis, as done for our human samples. T cells from stable plaques and the spleen were also included as comparators (**Figure 5A**). T cells were clustered and annotated as above (**Figure 5B & C**), identifying CD4 TN, CD4 TSM, CD4 T_H1_DM, CD4 T_reg_, CD8 TN, CD8 TSM, CD8^+^ T_c1-CTL_DM, proliferating T cells and a cluster of *Cd4^−^Cd8^−^* TSM cells in mice.

**Figure 5:**
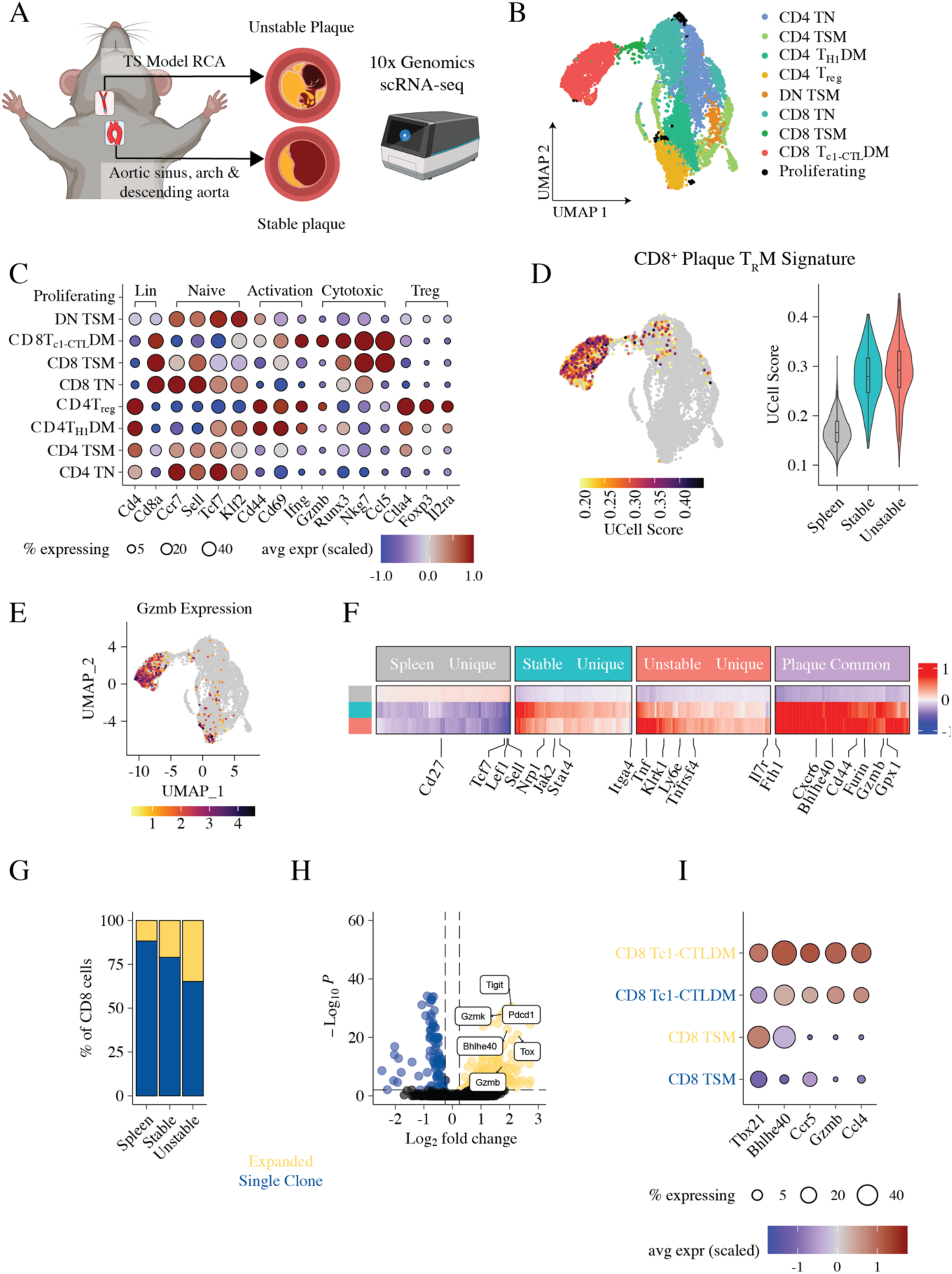
Clonally expanded, cytotoxic and T_R_M-like CD8^+^ T cells occupy murine unstable plaques. (A) T cells were flow cytometrically enriched from unstable (TS in the right carotid artery) and stable plaque tissue (aortic sinus, arch and descending aorta) from ApoE^−/-^ mice and then subjected to 10x genomics scRNA-sequencing with integrated αβ TCR sequencing. Matched splenic T cells were included as a peripheral comparator. (B) Annotated UMAP dimensionality reduction of T cell transcriptomes. (C) Select genes informing annotation. (D) CD8^+^ T_R_M signature (see Figure 4) projected onto CD8^+^ T cell clusters (left panel) and stratified by tissue (right panel). (E) *Gzmb* expression showing localisation predominantly to CD8^+^ T_c1-CTL_DM. (F) Heatmap of top 100 genes uniquely upregulated in T cells from each tissue, as well as those upregulated consistently in both plaque conditions versus the spleen. (G) Percentage of CD8^+^ T cells clonally expanded across tissues. (H) DEG analysis of expanded versus single-clone CD8^+^ T cells. (I) Expression of select genes from H by expanded and single-clone CD8^+^ TSM and CD8^+^ T_c1-CTL_DM.

The core plaque T_R_M signature that we identified in humans (**Figure 5D**) and established signatures (**Supplementary Figure 5B**) were all strongly expressed and upregulated in plaque versus splenic CD8^+^ T cells. Given that the transcriptional signatures of T_R_M in humans and mice can differ, we compared murine CD8^+^ T cells with T_R_M-high versus T_R_M-low signatures to define both species-specific and conserved features. We identified 111 genes that were consistently upregulated in both human and mouse T_R_M-high versus - low cells (e.g., *Cd69, Bhlhe40* and *Nr4a1-3*), as well as additional genes applicable specifically to mice (e.g., *Cxcr6;* **Supplementary Figure 5C & D**). Several genes associated with T_R_M in other tissues, including *Itga1* and *Itgae,* were not significantly upregulated, consistent with our human analyses. This further indicates that tissue-specific features govern T_R_M retention within atherosclerotic plaques.

Consistent with our findings in human atherosclerosis, T cells underwent major transcriptional remodelling in the plaque environment, with common and cluster-specific patterns of gene expression identified (**Supplementary Figure 5E & F**). Notably, CD8^+^ T_c1-CTL_DM cells (**Supplementary Figure 5G**) and cytotoxicity (e.g., *GZMB;* **Figure 5E** & **Supplementary Table 8**) were prominent in both unstable and stable plaques, with CD8^+^ T cells again being the main driver of plaque cytotoxic signatures (**Supplementary Figure 5F & H**). Moreover, we noted upregulation of additional genes consistent with enhanced cytotoxicity (e.g., *Klrk1, Tnf*)^85–87^ in unstable plaques (**Figure 5E**).

CD4^+^ and CD8^+^ T cells both demonstrated increased clonal expansion in plaques compared with the spleen. Moreover, expansion in both subsets was greater in unstable versus stable plaques (**Figure 5G & Supplementary Figure 6A**). As with humans, CD8^+^ T cells dominated the expanded pool in both tissues (**Supplementary Figure 6B**). This supports the conclusion that the CD8^+^ bias in humans is not the result of other factors (e.g., age or prior pathogen exposure) but reflects a fundamental characteristic of T cell immunity in atherosclerosis. We next compared expanded CD8^+^ T cells to single clones, noting enrichment for exhaustion-associated genes, including *TOX* and *PDCD1* (**Figure 5H**, **Supplementary Table 9**). CD8^+^ TX identified using reference mapping were found to be sparse peripherally, enriched within plaque tissue, and predominantly present within the CD8^+^ T_c1-CTL_DM cluster (**Supplementary Figure 6C**). In mice, unstable plaques contained more CD8^+^ TX than stable plaques (**Supplementary Figure 6D**). However, most importantly and paralleling our findings in human atherosclerosis, expanded CD8^+^ T cells in mice demonstrated significantly increased expression of cytotoxicity-associated genes (e.g., *Gzmb*) versus single clones, most notably expressed by expanded CD8^+^ T_c1-CTL_DM (**Figure 5H & I**).

Overall, these data confirm that the key T cell states we identified in human atherosclerosis are reflected in unstable and stable plaques within the TS model. They also further enabled the identification of putative species-conserved regulators of plaque CD8^+^ T cell tissue residency. Moreover, we confirm that the accumulation of clonally expanded CD8^+^ T cells with highly cytotoxic features is a core feature of unstable atherosclerotic plaques.

### TCR repertoire diversity is required for CD8^+^ T cells to drive plaque instability

Our data support the model that antigen-driven CD8^+^ T cell responses are a fundamental driver of plaque instability. To mechanistically test this idea, we performed experiments in OTI mice, which have a restricted CD8^+^ TCR repertoire where the vast majority of CD8^+^ T cells are specific for ovalbumin, a foreign antigen they are not exposed to.^88^ These mice, therefore, have limited capacity to respond to atherosclerosis-related antigens.

OTI mice rendered athero-susceptible via a single AAV8-PCSK9 injection^9,51,52^ and then subjected to TS surgery showed significant reductions in unstable and stable atherosclerosis development compared with wild-type (WT) control mice (**Figure 6A–C**). Focusing on unstable atherosclerotic disease in the right carotid artery of TS mice, the absolute CD68^+^ area was reduced in OTI versus WT mice, reflecting reduced foam-cell development (**Figure 6D & E**). This was accompanied by significant reductions in the number of CD68^+^ (**Figure 4F & G**) and CD68^+^MHCII^+^ (**Figure 6H**) macrophages, and the near-complete absence of Ly6G^+^ neutrophils (**Figure 6I**) in OTI mice. Plaques from OTI mice also had significantly increased nucleus density, consistent with reduced necrotic acellular area and thus increased plaque stability (**Figure 6J**). Finally, OTI mice did not develop intraplaque haemorrhage (IPH), a fundamental feature of plaque instability,^53,54^ which was present in ∼50% of WT mice as expected^26^ (**Figure 6K & L**). Combined, these data demonstrate that antigen-driven CD8^+^ T cell responses are a critical driver of atherosclerotic plaque destabilisation.

**Figure 6:**
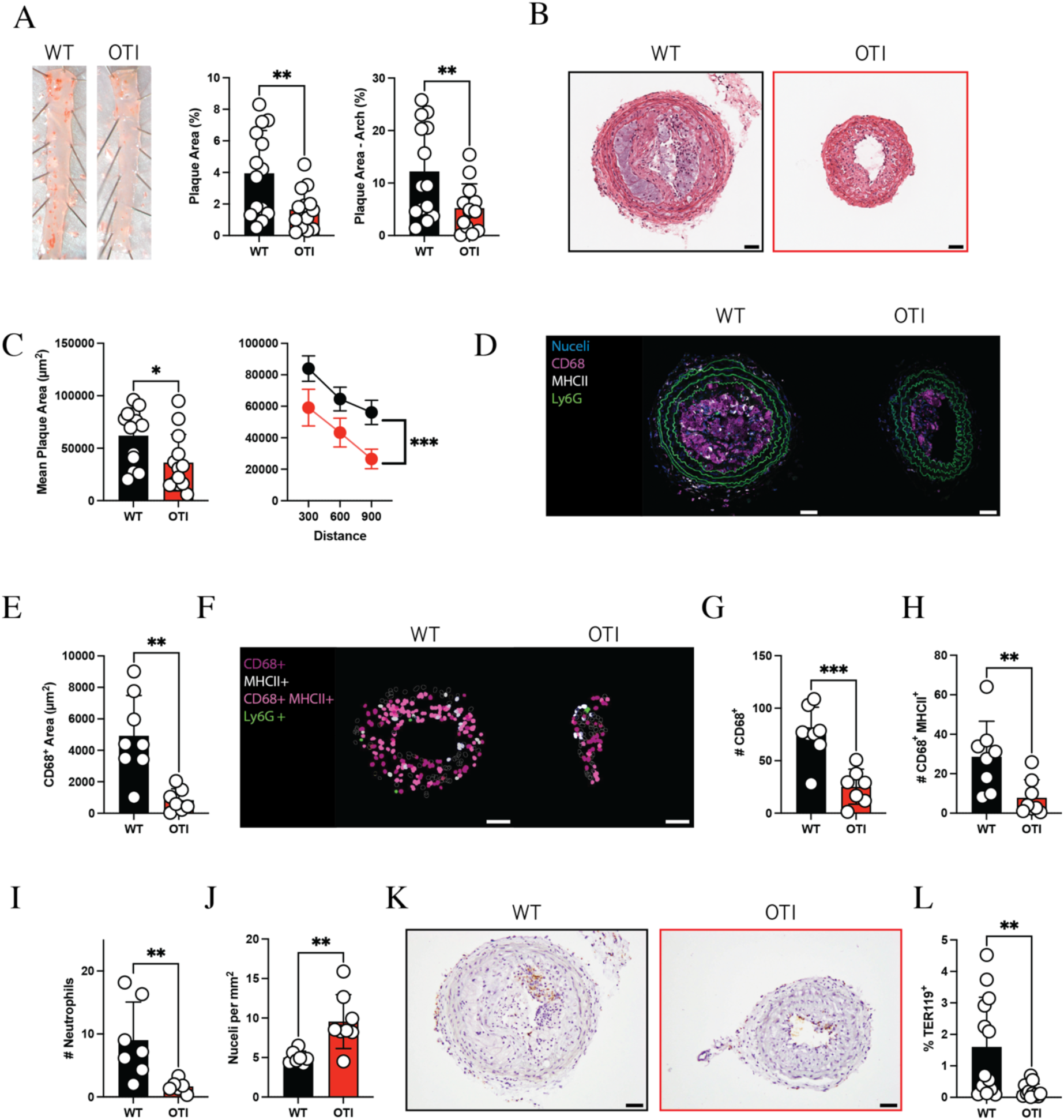
CD8^+^ TCR diversity is required to drive plaque instability. C57BL/6 (WT) and TCR transgenic OTI mice received a single injection of AAV8-PCSK9 to render them athero-susceptible before being fed a high-fat diet and then subjected to TS surgery. All analyses were conducted at 7 weeks post TS surgery, comparing WT versus OTI mice. (A) Representative images of Sudan IV *en face* plaque staining of stable disease in the aorta (left panel), along with quantification of % plaque area of the entire vessel (middle panel) and aortic arch (right panel). (B) Representative H&E images of the right carotid artery containing unstable plaques from WT and OTI mice. (C) Assessment of unstable plaque area quantified 300, 600 and 900 μm distal from the TS. (D) Representative immunofluorescence images of CD68, MHCII and Ly6G expression. (E) CD68^+^ plaque area. (F) Representative StarDist images and quantification of (G) macrophages (CD68^+^), (H) MHCII^+^ macrophages, (I) neutrophils (Ly6G^+^) and (J) nucleus density. (K) Representative images and (L) quantification of IPH via TER119 staining. In A–C & L, *n =* 13–14 (WT) & 13–14 (OTI). In E–J, *n* = 6–8 (WT) & 6–8 (OTI). Scale bars = 50 μm. All bar plots are presented as mean +/-SD except for the line graph in C (right panel), which is mean +/− SEM. Data were analysed using Welch’s t test (A, E, I, J), Student’s t test (C; left panel, G), Mann–Whitney test (H & L) and ordinary two-way ANOVA (C; right panel). *\*p<0.05; **p<0.01; ***p<0.001*.

### CD8^+^ T cell cytotoxicity correlates with carotid plaque instability and predicts clinical outcomes in patients with atherosclerosis

We next tested in a large patient population our hypothesis that CD8^+^ T cells associate with plaque instability. To do this, we leveraged the AtheroExpress Biobank, which uniquely combines histological and molecular analysis of human carotid plaques with three years of clinical follow-up for over 1000 patients.^89,90^ In all analyses, we considered key confounders (age, gender, hypertension, diabetes, smoking, lipid-lowering drugs, antiplatelet medication, eGFR and BMI).

We first tested whether key transcriptional features of cytotoxic CD8^+^ T cell responses (assessed via bulk RNA-sequencing) correlated with histologically defined plaque instability in the carotid plaques’ of 1093 patients. We selected *GZMB* (prototypical cytotoxic protein), *PRDM1* and *RUNX3* (both transcriptional regulators of CD8^+^ T cell cytotoxic function) as they were all highly expressed by plaque CD8^+^ T cells (**Figure 7A**). Moreover, *PRDM1*^+^*RUNX3*^+^ CD8^+^ T cells demonstrated significantly upregulated cytotoxic gene expression versus all other CD8^+^ T cells (**Figure 7B**). Notably, these drivers of cytotoxicity were all significantly associated with one or more features of instability (**Figure 7C & Supplementary Table 10**). *GZMB* alone, for example, was significantly associated with reduced smooth muscle cell content, increased lipid content and increased IPH.

**Figure 7:**
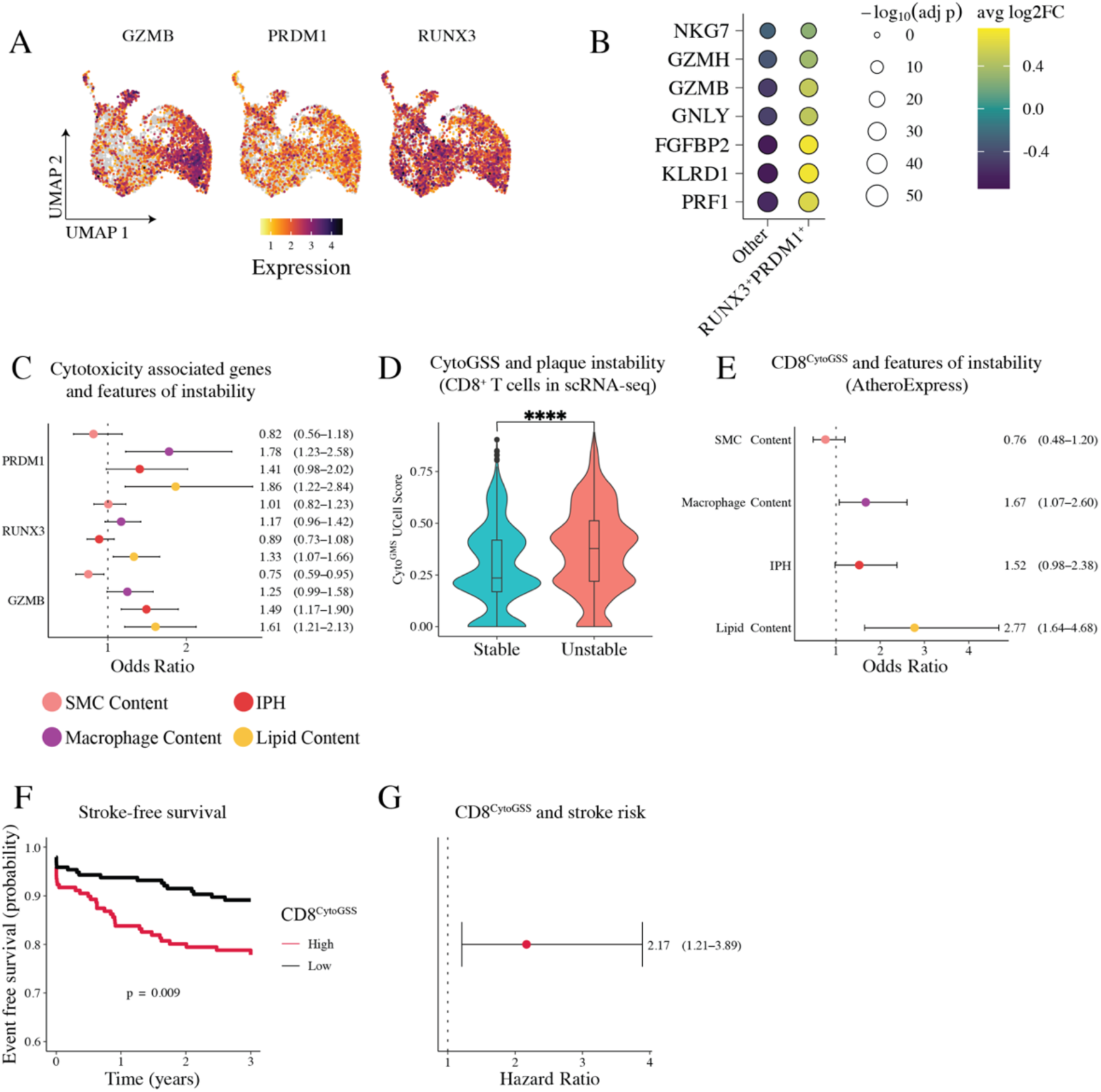
Intraplaque CD8^+^ T cell cytotoxicity is associates with plaque instability and predicts recurrent events in patients with atherosclerosis. (A) Analysis of *GZMB*, *PRDM1* and *RUNX3* expression in atherosclerotic plaque CD8^+^ T cells assessed by scRNA-seq. (B) Expression of cytotoxicity-associated genes in *RUNX3*^+^*PRDM1*^+^ versus all other CD8^+^ T cells in atherosclerotic plaques in AtheroExpress. (C) Association between *PRDM1*, *RUNX3* and *GZMB* and key features of plaque instability in AtheroExpress. (D) CytoGSS gene-set scoring (*GZMB*, *RUNX3* and *PRDM1*) in unstable versus stable plaque CD8^+^ T cells assessed by scRNA-seq. (E) Association between CD8^CytoGSS^ (CytoGSS + *CD3E & CD8B*) and features of plaque instability in AtheroExpress. (F) Stroke-free survival in patients with high versus low CD8^CytoGMS^ in AtheroExpress. (G) Estimated future stroke risk in patients with high versus low CD8^CytoGMS^ in AtheroExpress. IPH = intraplaque haemorrhage; SMC = smooth muscle cell. *n* = 1093 patients in dataset. Confidence intervals are shown for data in C, E and G. All AtheroExpress analyses are adjusted for age, sex, year of surgery, hypertension, diabetes, smoking, LDL-C level, lipid-lowering drugs, antiplatelet drugs, eGFR, BMI, history of CVD and level of stenosis. Data were analysed using a Mann–Whitney test (D); *\*\*\*\*p<0.0001*.

We next utilised gene-set scoring, combining *GZMB, PRDM1* and *RUNX3* into a cytotoxic signature, which we first validated was significantly upregulated in unstable versus stable plaque CD8^+^ T cells in our scRNA data (**Figure 7D**). To link cytotoxicity directly to CD8^+^ T cells in AtheroExpress, we additionally incorporated *CD3E* and *CD8B* into this signature (CD8^CytoGSS^). Our analysis revealed CD8^CytoGSS^ was significantly associated with features of plaque instability including increased lipid content and increased macrophage content, with a strong trend towards increased IPH (**Figure 7E & Supplementary Table 10**).

A major advantage of the AtheroExpress Biobank is the unique opportunity to assess the longitudinal relationship between patients’ intraplaque gene expression at presentation and clinical outcomes over a three-year follow-up. As plaques were isolated from carotid arteries, we focused on stroke as the primary major outcome. Vitally, we discovered that patients with a high CD8^+^ T cell cytotoxicity score had a significantly increased risk of stroke (hazard ratio 2.17, 95% CI 1.21-3.89; **Figure 7F & G, Supplementary Table 11**). Together, these results demonstrate that cytotoxic CD8⁺ T cell activity within atherosclerotic plaques is not only a hallmark of instability, but also a predictor of future events, underscoring the mechanistic importance and potential as a clinically actionable biomarker of high-risk disease.

## Discussion

Here, we leveraged well validated approaches (NIRAF imaging^91,92^ and our robust TS mouse model^83,84,93^) to precisely dissect T cell responses in unstable versus stable human and mouse atherosclerosis. Through comprehensive analysis of single-cell gene expression, T cell receptor repertoires (i.e., clonality), mechanistic animal modelling and interrogation of a large prospective cohort study, we discovered that autoimmune-like cytotoxic CD8^+^ T cell responses are a central driver of atherosclerotic plaque instability and increased risk of major adverse cardiovascular events. Against the backdrop of emerging efforts to therapeutically target adaptive immunity in cardiovascular disease,^33,94,95^ our findings position CD8⁺ T cell immunity as a tractable and timely target with both prognostic value and therapeutic potential, offering opportunities for improved risk stratification and the development or repurposing of immunomodulatory strategies to stabilise plaques and prevent cardiovascular events.

A central novelty is our discovery that T cell responses and their clonality differ significantly in precisely stratified unstable compared to stable plaque tissues. Our approaches (NIRAF and TS) removed limitations inherent to comparing, for example, clinically or histologically defined disease stages, as such plaques will still contain stable and unstable regions, potentially confounding results. Beyond our own discoveries, discussed further below, these data provide a foundational resource and framework within which additional drivers of instability can be explored.

An important aspect of this work was exploring the interaction between clonal expansion, T cell activation states and instability. Existing data supported that T cells are generally clonally expanded in atherosclerotic plaques.^9,13,14,46^ Nevertheless, we were conscious of technical and statistical limitations in the foundational data from carotid plaques (described in results above). Our study importantly clarifies that T cells demonstrate the reduced TCR repertoire diversity consistent with greater expansion in plaques than blood. Moreover, we discovered that TCR repertoires are significantly more similar in patients’ plaques than blood, and that repertoires differ markedly between unstable and stable plaques. These data thus provide two key insights. First, similar or identical antigens likely drive atherosclerosis-associated T cell responses across patients. Second, unique antigen environments could underpin the activation of T cells with specificities uniquely capable of driving instability.

When analysing both T cell transcriptional signatures and clonality in unstable versus stable plaques, CD8^+^ T cells stood out. For example, CD8^+^ T cells were overall the most expanded population in both human and mouse plaques. As the atherosclerotic mice were relatively young, these data demonstrate that this CD8^+^-biased response is not a ‘side-effect’ of factors like ageing known to impact T cell responses^19,96^, but a fundamental characteristic of atherosclerosis. Most critically, unstable plaques were markedly enriched for cytotoxic CD8^+^ T cell signatures. This heightened cytotoxic profile was intrinsically linked to clonally expanded CD8^+^ T cells, which demonstrated the greatest cytotoxic gene expression. Thus, we discovered a compelling connection between CD8^+^ T cell clonal expansion, cytotoxicity and atherosclerotic plaque instability. This suggests that antigen-driven autoimmune(-like) cytotoxic CD8^+^ T cells drive plaque instability.

Another point of novelty is our significantly improved understanding of exhausted (TX) and tissue-resident memory (T_R_M) CD8^+^ T cells in plaques. Both of these T cell states are dependent on antigen-driven activation and have been inferred to exist in human plaques based on the expression of a restricted number of gene/protein markers.^9,50,72,74^ However, detailed understanding has been limited, with even their abundance in plaques being unclear. First, we identified that exhaustion is predominantly a feature of CD8^+^ (vs. CD4^+^) T cells in atherosclerosis, that CD8^+^ TX are enriched in plaques, and that exhaustion signatures correlate with TCR signalling. Overall, the presence of CD8^+^ TX and their apparent reliance on TCR signalling provide further support to the hypothesis that chronic T cell–mediated inflammation within plaques is driven by antigen-dependent and likely autoimmune mechanisms.

With respect to tissue residency, we surprisingly identified that genes previously used to identify T_R_M in plaques (e.g., *ITGAE, ITGA1,* ZNF683), which are important in other tissues, do not appear strongly relevant to residency in plaques. Thus, we then explicitly defined the transcriptional identity of T_R_M cells in human and mouse plaques, including their increased expression of *RGCC, BHLHE40, CD69, TFRC* and *ICOS*. Our identification of this “plaque-specific” T_R_M signature is consistent with tissue-dependent adaptations noted in many other studies of numerous human and mouse tissues.^67,68,75–77^ Critically, our approach enabled us to finally quantify T_R_M in plaques. In contrast to prior studies,^9,72^ we now clarify that the majority of CD8^+^ T cells within plaques are tissue resident. This fundamentally changes our perception, as previously it would have been interpreted that most plaque T cells are transient “visitors” to the plaque.

Interestingly, we also discovered that CD8^+^ T_R_M are enriched for exhaustion-associated genes, including *TOX,* in plaques versus other healthy human tissues. While T_R_M and TX differentiation was previously considered independent, our data aligns with recent studies evidencing that these states can converge in chronically inflamed tissue.^65,81,82^ Thus, we extend this new paradigm that T_R_M can become exhausted to the atherosclerosis setting.

Considering our data overall, cytotoxicity provides a logical mechanism by which CD8^+^ T cells could promote plaque destabilisation and rupture through killing cells integral to maintaining the protective fibrous cap (e.g., smooth muscle cells)^40,41^ or increasing the prevalence of pro-atherogenic dead/dying cells. However, CD8^+^ T cell–mediated killing of pathogenic cells could also be protective.^35,36,42^ Thus, the antigen specificity of CD8^+^ T cells is likely integral to their overall impact. To address the net contribution of antigen-driven CD8^+^ T cell responses in plaque destabilisation we combined TCR-transgenic mice, AAV8-PCSK9-induced atherosclerosis and our TS model. In doing so, we explicitly demonstrate for the first time that endogenous antigen-mediated CD8^+^ T cell activation is critical to the development of atherosclerosis and, more importantly, the destabilisation of plaques towards rupture. Moreover, given mice in these experiments were housed in the relative absence of pathogens, these data also further support that the responses driving instability are directed against self-antigens and are thus autoimmune.

Of utmost importance, and as a final step in our study, we confirmed the clinical relevance of our findings leveraging the unique AtheroExpress Biobank. Here, we discovered that signatures of CD8^+^ T cell cytotoxicity in excised atherosclerotic plaques correlated significantly with histologically defined instability in >1000 patients. Moreover, this cytotoxic CD8^+^ signature was directly associated with increased risk of subsequent stroke. Thus, these data in a large prospective patient cohort support that CD8^+^ T cell responses drive instability to directly cause major adverse cardiovascular events. This is the first data robustly correlating CD8^+^ T cell responses within plaques with increased risk of cardiovascular events.

## Conclusions

This study is the first to assess T cell immunity explicitly in unstable compared to stable atherosclerotic plaque in both humans and mice. We identified compelling cellular, molecular and mechanistic evidence that autoimmune(-like) cytotoxic CD8^+^ T cell responses drive plaque instability. Of central clinical relevance, we discovered that cytotoxic CD8^+^ T cell signatures in carotid atherosclerosis are significantly and prospectively associated with the incidence of stroke, suggesting prognostic utility for identifying patients at high risk of secondary events. Moreover, the growing arsenal of T cell–centric therapeutics available for numerous non-cardiovascular indications and emerging preclinical successes in the cardiovascular setting make T cell modulation a highly viable and promising treatment option for atherosclerosis.^33,35,95,97,98^ Our data thus support leveraging the critical mass of expertise and cutting-edge therapeutic platforms to prevent autoimmune(-like) CD8^+^ T cell responses as a potentially pivotal driver of plaque rupture and major adverse cardiovascular events.

## Supplementary Figures

**Supplementary Figure 1:**
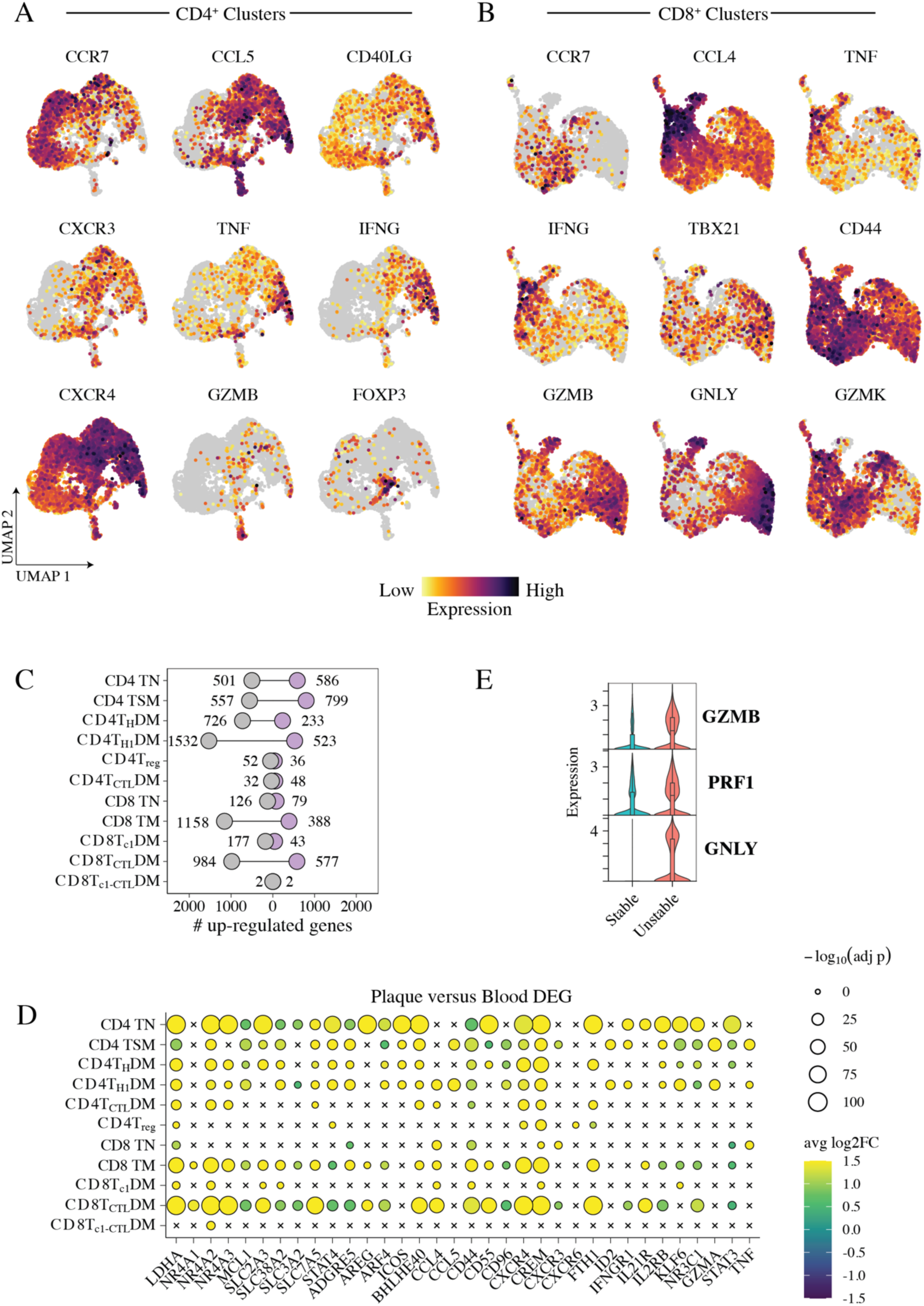
Distinct T cell signatures in human plaques versus blood and increased cytotoxic profiles in unstable versus stable plaques. These data complement and expand on Figure 1. Carotid plaques were isolated from patients undergoing endarterectomy and, using NIRAF imaging, classified into unstable (NIRAF^High^) and stable (NIRAF^Low^) atherosclerotic plaque. T cells were flow cytometrically enriched from unstable and stable segments and then subjected to 10x genomics scRNA-sequencing. Matched peripheral blood T cells were also included. Expression of key genes informing annotation in (A) CD4^+^ and (B) CD8^+^ T cell clusters. (C) Bubble plot presenting the number of genes differentially expressed by each cluster in plaque- versus blood-derived T cells and (D) the upregulation of select genes from this analysis. (E) Expression of key cytotoxicity-associated genes in unstable versus stable plaque-derived T cells.

**Supplementary Figure 2:**
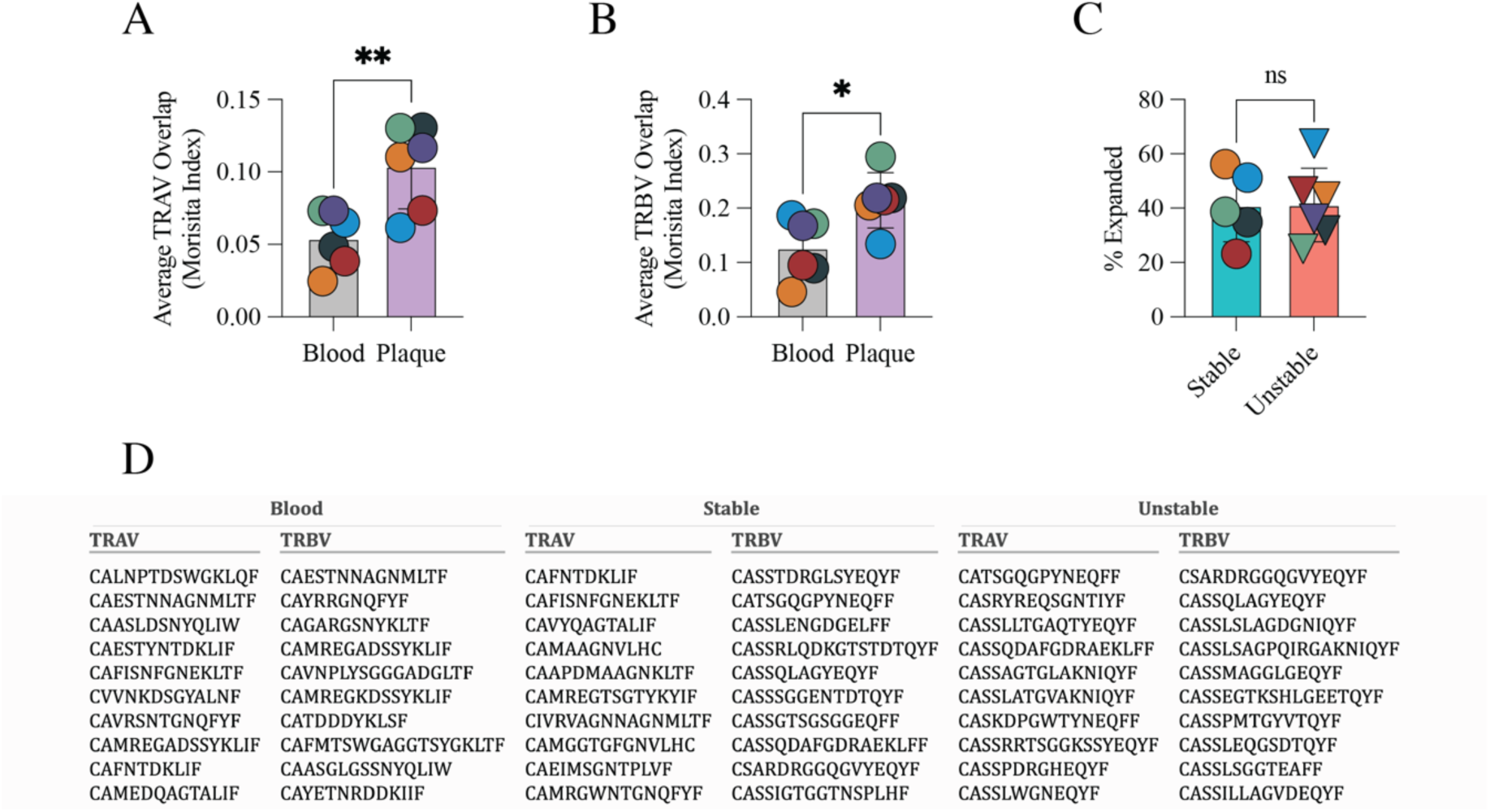
T cell clonality and TCR repertoires in human atherosclerosis. These data complement and expand on Figure 2. Morisita index (average index for each patient vs. all others) comparing (A) TCRα and (B) TCRβ chain similarity in patient plaques versus blood. (C) Percentages of T cells clonally expanded in unstable versus stable plaque T cells. (D) T cell receptor sequences for the top 10 T cell clones in unstable versus stable plaques and versus blood.

**Supplementary Figure 3:**
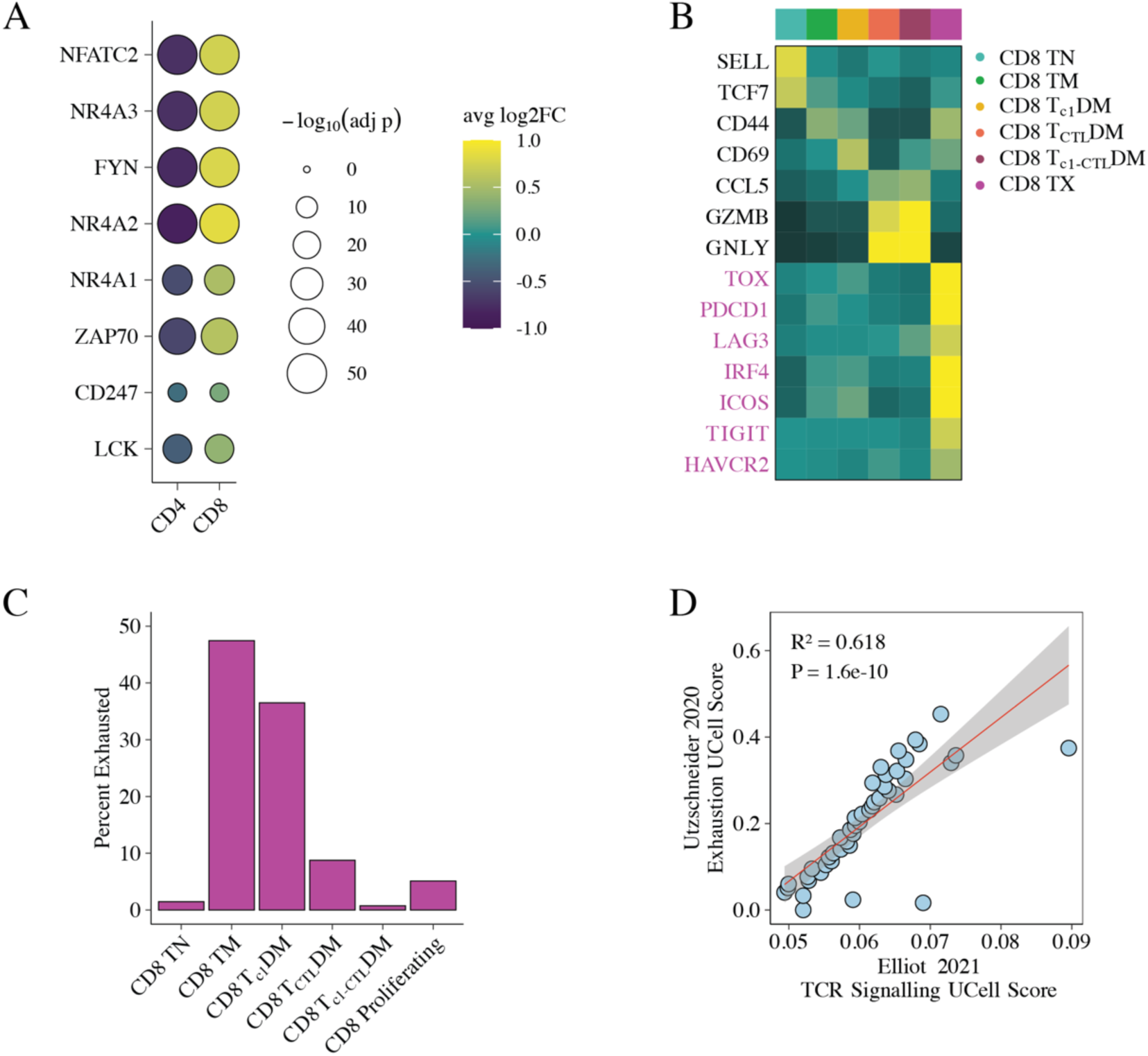
T cell exhaustion in human atherosclerosis. These data complement and expand on Figure 3. (A) Expression of genes indicative of T cell receptor signalling in CD4^+^ versus CD8^+^ plaque-derived T cells. (B) Heatmap of select genes, including exhaustion-associated genes (highlighted in purple), in each CD8^+^ T cell cluster versus cells identified as TX. (C) Percentages of exhausted cells originating from each CD8^+^ T cell cluster based on manual annotation. (D) Correlation between T cell exhaustion gene-set scoring based on Utzschneider *et al*. and TCR signalling based on Elliot *et al*., excluding all genes present in both gene sets.

**Supplementary Figure 4:**
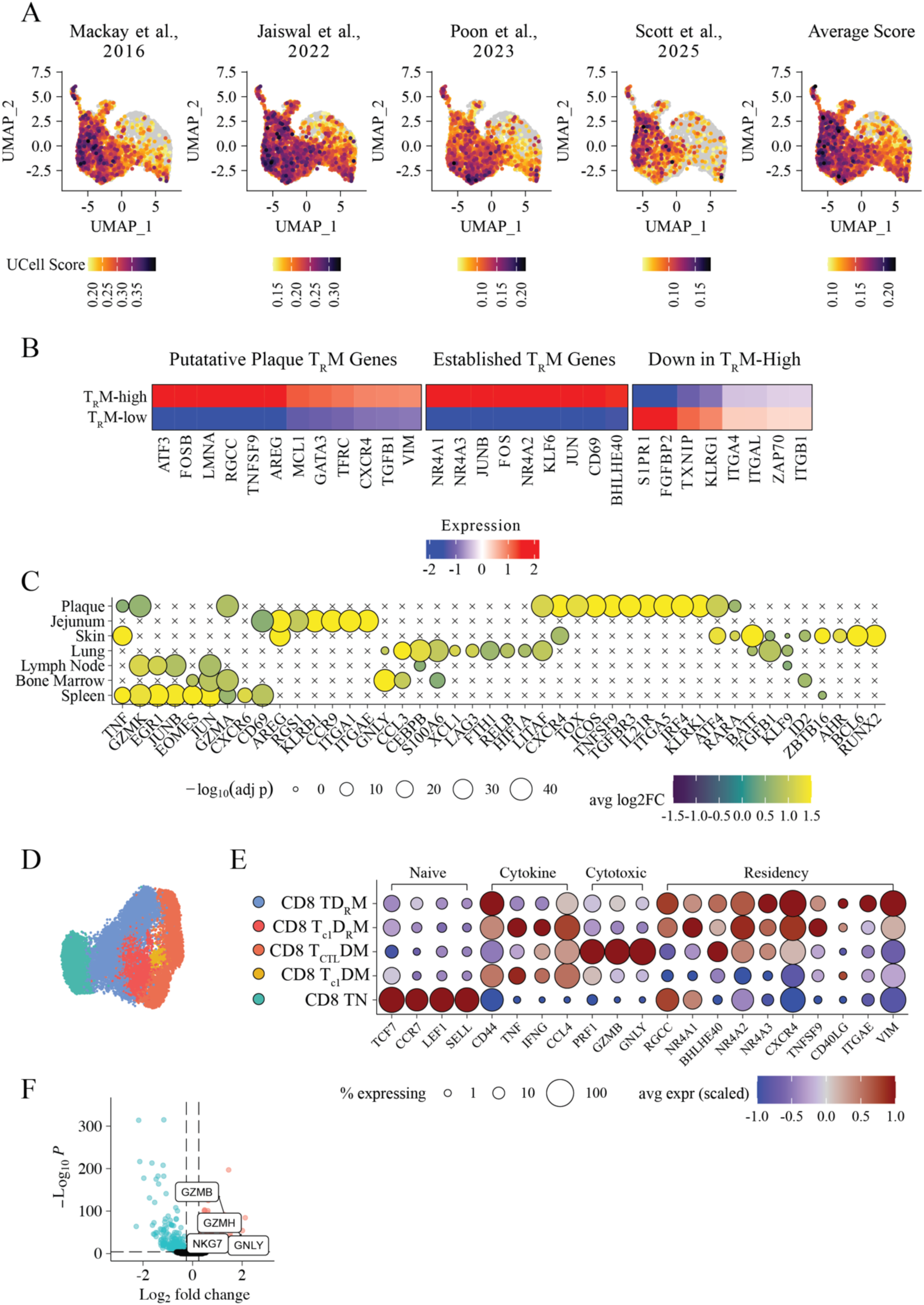
Defining the transcriptional identify of CD8^+^ tissue-resident T cells in in human atherosclerotic plaques. These data complement and expand on Figure 4. (A) Feature plot of UCell scores from established tissue-residency gene sets from multiple published sources. (B) Select genes differentially expressed in CD8^+^ T_R_M-high versus T_R_M-low signature cells. Genes consistently reported in other studies versus those that may be of increased relevance to plaque CD8^+^ T cells are indicated. (C) We integrated our plaque data with multi-tissue T cell scRNA-seq data from Poon *et al.* to compare plaque T_R_M to those in other well-characterised tissues. Select genes differentially expressed by T_R_M-high cells in plaque CD8^+^ T cells versus multiple other human tissues. (D) UMAP of CD8^+^ T cell clusters in our integrated dataset and (E) the select genes used to annotate these. (F) Volcano plot of genes differentially expressed between T_R_M-high cells in unstable versus stable plaques, highlighting key cytotoxic genes.

**Supplementary Figure 5:**
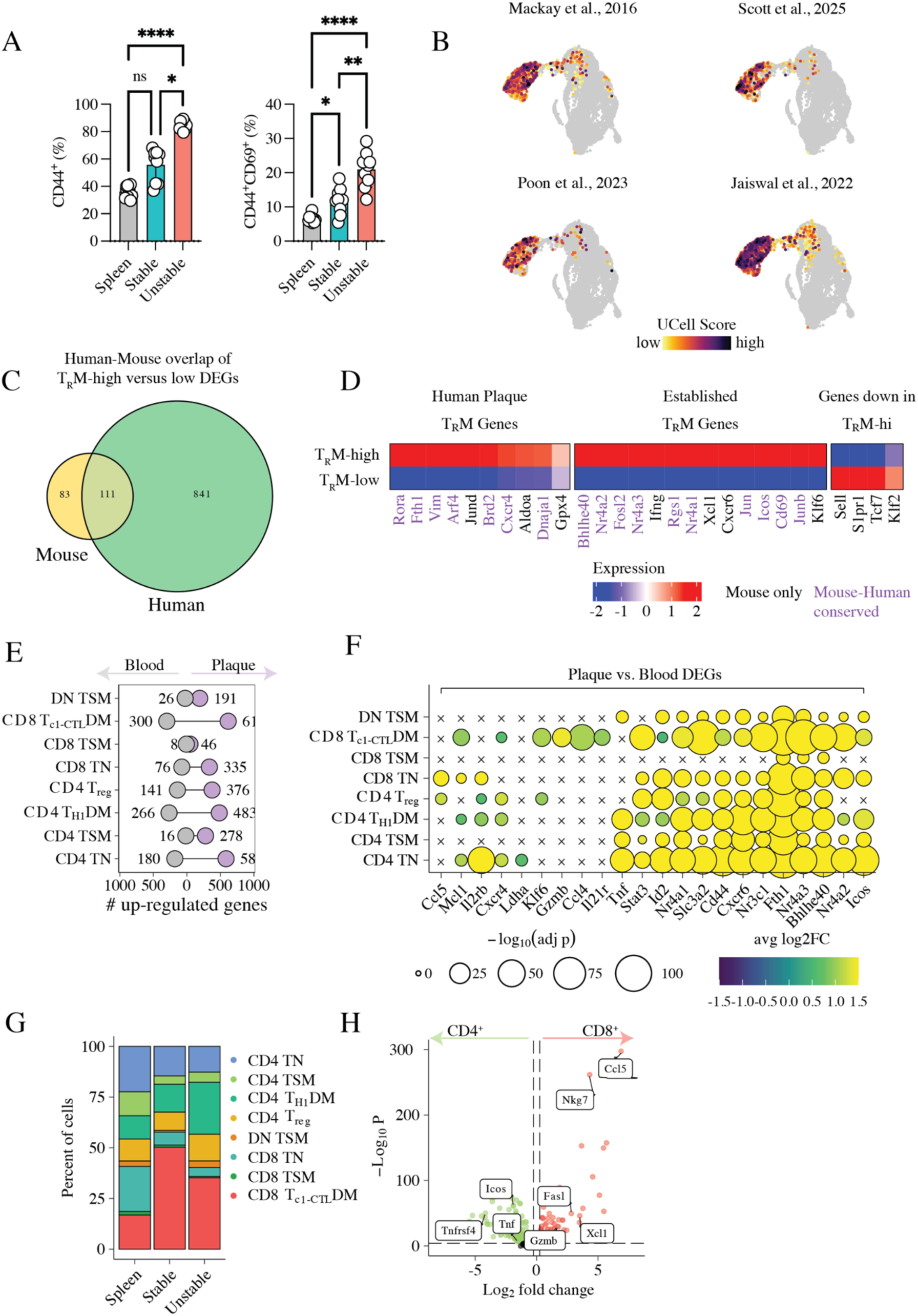
CD8^+^ T cell activation states identified in human plaques are reflected in plaques from the TS mouse model of plaque instability. These data complement and expand on Figure 5. (A) CD3^+^ T cells were assessed for CD44 and CD69 expression using flow cytometry to quantify their activation status in unstable (*n* = 9) versus stable (*n* = 9) plaques, including the spleens (*n* = 10), as a peripheral comparator, as in scRNA analyses. Left panel was analysed using Kruskal–Wallis with Dunn’s multiple comparisons test; right panel was analysed using Brown–Forsythe and Welch ANOVA with Dunnett’s T3 multiple comparisons test. \**p<0.05*; \*\**p<0.01*; \*\*\*\**p<0.0001*; ns = not significant. All subsequent analyses leveraged our scRNA-seq data or murine atherosclerosis in the TS model. (B) Feature plot of UCell scores from established tissue-residency gene sets from multiple published sources projected onto CD8^+^ T cell clusters. (C) Venn diagram of the number of genes upregulated in human and mouse T_R_M-high versus T_R_M-low CD8^+^ T cells and their overlap. (D) Select genes differentially expressed in CD8^+^ T_R_M-high versus T_R_M-low signature cells. Genes consistently reported in other studies versus those that may be of increased relevance to plaque CD8^+^ T cells are indicated. Genes upregulated in both human and mouse T_R_M-high cells are highlighted in purple. (E) The numbers of genes differentially expressed in each T cell cluster between plaque (purple) and splenic (grey) T cells and (F) the upregulation of select genes from this analysis. (G) Proportions of each T cell cluster in blood, stable and unstable plaques. (H) Volcano plot of differentially expressed genes between CD8^+^ and CD4^+^ plaque T cells.

**Supplementary Figure 6:**
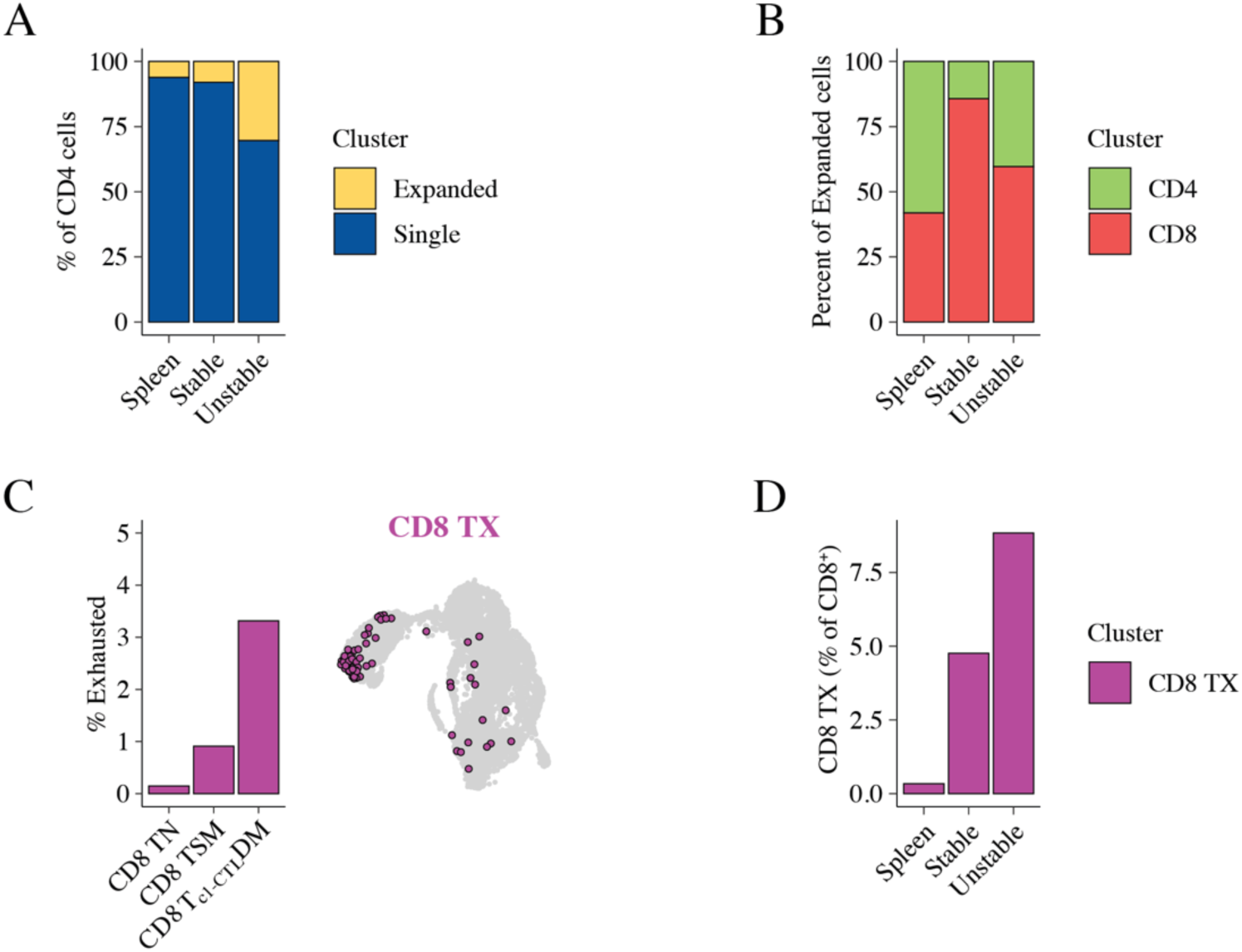
T cell expansion and exhaustion in the TS mouse model of plaque instability. These data complement and expand on Figure 5. All data are from scRNA-seq analysis. (A) Percentage of CD4^+^ T cells clonally expanded across tissues. (B) Percentages of expanded cells in each tissue that were CD4^+^ versus CD8^+^. (C) CD8^+^ TX were identified using reference mapping. Percentage of TX coming from each CD8^+^ T cell cluster (left panel) and positioning of TX on the cluster UMAP. (D) Percentages of CD8^+^ TX cells in plaque tissues versus blood based.

## Methods

### Animals

All animal procedures were approved by the Animal Ethics Committee of the Alfred Medical Research and Education Precinct (AMREP), Melbourne, Australia. C57BL/6J, ApoE^−/-^ and OTI mice were bred and maintained at the AMREP Animal Centre with constant access to food and water.

### Tandem stenosis surgery

The TS surgery has been extensively described previously.^83^ Briefly, 6- to 12-week-old mice (ApoE^−/-^, C57BL/6 or OTI) were fed a high-fat diet (HFD; SF00–219, 22% fat and 0.15% cholesterol, Specialty Feeds) for 6 weeks. Mice were then anaesthetised using inhaled 4% isoflurane. Carprofen (5 mg/kg, Lyppard, Keysborough, VIC, Australia) was administered subcutaneously as a painkiller. Bupivacaine (2 mg/kg) was given subcutaneously before an incision was made in the neck and the right common carotid artery was exposed. A TS with 150 μm outer diameter was then introduced, with the distal point 1 mm from the carotid artery bifurcation and the proximal point 3 mm from the distal stenosis. To achieve the stenosis diameter, a 6–0 TI-CRON blue braided-polyester fibre suture was fed underneath the carotid artery. A 150 μm needle was then laid on top of the artery and the suture tied over the needle. The needle was then removed. Animals were then recovered and maintained on HFD. At end points, mice were euthanised, blood was collected by cardiac puncture if required and then mice were perfused through the left ventricle using 20 mL of PBS. Tissues were then collected for analyses.

### Induction of atherosclerosis using AAV8/mPCSK9

To induce atherosclerosis in OTI and C57BL/6J animals, they received a single intravenous injection of the adeno-associated virus (AVV) AAV8/D377Y-mPCSK9.^99^ Each mouse received 1×10^11^ vg of AAV one week prior to enrolment in the TS surgery protocol (described above). Animals were maintained on an HFD diet for 7 weeks post TS surgery and then euthanised for analysis. LDL-C levels were measured in these mice at end point and LDLR downregulation in the liver was confirmed by Western blot. Only mice with LDLR downregulation and hypercholesterolemia (plasma LDL-C ≥250 mg/dL) were included in analyses.

### Isolation of single-cell suspensions from humans and mice

To isolate peripheral blood mononuclear cells (PBMCs) from human blood samples, blood was collected in BD Vacutainers (lithium heparin tubes) via an arterial line. Blood was diluted 1:2 in HBSS and separated by layering on Ficoll-Paque Plus density separation media followed by centrifugation (400 xg, 40 min at room temperature with no break). The PBMC layer was then retrieved. Mouse spleens were homogenised through a 70 μm filter, with the resulting solution then filtered through a 40 μm filter. To isolate single-cell suspensions from murine vessels and human endarterectomy samples, the tissues were finely minced and then placed into ice-cold 0.9 mM CaCl2 (in PBS) containing Liberase TL (0.4 U/mL) and DNase1 (50 U/mL). These were then transferred into a 37 °C water bath and triturated using a 1 mL pipette (set to 750 μL) every 15 min for a total of 45 min. Samples were then filtered through a 70 μm filter, with manual homogenisation also applied using a syringe plunger to disaggregate any remaining tissue. Samples were then refiltered through a 40 μm filter, providing a single-cell suspension.

### Flow cytometry

Single-cell suspensions were centrifuged and then resuspended in fixable viability dye for 15 min. Following washing in PBS and centrifugation, cells were stained with a master mix of fluorochrome-conjugated antibodies for 30 min at 4 °C. All steps were conducted at 4 °C prior to acquisition. Centrifugation was always 400 xg for 5 min. Data were acquired from stained cells using a BD LSR Fortessa II (BD Bioscience, USA). Data analysis was performed using FlowJo (Flowjo LLC, USA).

### Single-cell RNA sequencing – mouse

To analyse cells from unstable plaques, right carotid arteries were isolated from ApoE mice 4 and 9 weeks post TS surgery to ensure representation of cells across the instability spectrum.^83^ To assess stable plaque cells, the aortic sinus, arch and descending aorta were isolated from identically treated mice that did not undergo TS surgery. All vascular tissue was then cleaned of perivascular fat and digested (as described above) to obtain single-cell suspensions. Splenic single-cell suspensions were also obtained from the same mice. To obtain sufficient cells, we pooled vessels from 8–12 mice per group in two independent experiments. Cells were then stained with oligonucleotide-labelled antibodies (BioLegend TotalSeq-C; protein labelling/hashing) and fluorescently labelled antibodies alongside fixable viability dye and calcein-AM to allow for the sorting of live metabolically active T cells. T cells were then FACS sorted and subject to scRNA-seq with matched TCR sequencing using the 10x Genomics Chromium and 5’ Immune Profiling (V1.1) Kits as per the manufacturer’s instructions. Sort purification was performed on a BD FACSAria Fusion or Aria. Sequencing was performed on a NovaSeq6000.

### Human specimens and patient characteristics

Six patients (**Supplementary Table 12**) undergoing carotid endarterectomy at the Alfred hospital in Melbourne, Australia, were enrolled in a study approved by the Alfred Research Alliance Human Research Ethics Committee (Project ID: 580/20). Written consent was obtained.

### Near-infrared autofluorescence (NIRAF) imaging

Endarterectomies were first assessed for NIRAF as described previously^91^ using an IVIS Lumina III imaging system (PerkinElmer, Shelton, CT, USA). Briefly, a visible-light photograph was taken to provide an anatomical underlay. NIRAF images were then obtained using two channels (excitation/emission 640/710 nm and excitation/emission 740/790 nm). Areas of high (unstable plaques) versus low (stable plaques) relative NIRAF were then macroscopically dissected and separated prior to processing and single-cell RNA-seq.

### Single-cell RNA sequencing – human

Carotid endarterectomies were separated into unstable and stable regions using NIRAF and then single-cell suspensions were prepared. PBMCs were also isolated from matched blood samples. Cells were then stained with oligonucleotide-labelled antibodies (BioLegend TotalSeq-C; protein labelling/hashing) and fluorescently labelled antibodies alongside fixable viability dye and calcein-AM to allow for the sorting of live metabolically active T cells. T cells were then FACS sorted and subjected to scRNA-seq with matched TCR sequencing using the 10x Genomics Chromium and 5’ Immune Profiling (V2) Kits as per the manufacturer’s instructions. Sort purification was performed on a BD FACSAria Fusion or Aria. Sequencing was performed on a NovaSeq6000.

### Processing and analysis of scRNA-seq data

Data were initially processed on CellRanger and then analysed predominantly using Seurat^100^ and scRepertoire^48,101,102^ architecture. Briefly, raw data were cleaned using SoupX. Demultiplexing of hashed samples was performed using MULTIseqDemux. TCR data were added to single-cell objects using scRepertoire. Data were then integrated using scTransform (in-house data) or FastMNN (when combining with publicly available data). scGate and PorjectTILs were used to isolate or reference-map T cell subsets as detailed in the text. For TCR analysis, clonal expansion was calculated based on the same TCR being detected on ≥2 cells (for mouse data this was across all cells; for humans this was within each patient) versus once (i.e., a single clone). To perform GRN analysis, the SCENIC pipeline^64^ was used. For differential testing, FindMarkers (pairwise comparison) or FindAllMarkers (multiple comparisons) functions were used implementing the MAST model. Only genes with a Bonferroni corrected p value <0.05 were considered statistically different.

### Histological analysis of murine atherosclerotic plaques

After perfusion, the entire aortic arch with the brachiocephalic artery was collected and transferred into 10% NBF for Sudan IV *en face* plaque-staining analyses. The aortic sinus and right and left carotid arteries were collected and fixed in 4% paraformaldehyde (PFA) for 3 h on ice and then incubated in 20% sucrose overnight at 4 °C. Samples were embedded upright in optimal cutting temperature (OCT) compound (Sakura Finetechnical, Torrance, CA, USA), snap-frozen in liquid nitrogen and stored at −80 °C for subsequent histological analysis. 6 μm thick cryosections of the aortic sinus and carotids were prepared using a cryostat (Leica CM 1950). Sections were histologically stained with standard Mayer’s haematoxylin/eosin (H&E) and imaged using an Aperio Scanscope AT Turpo. Quantification of carotid histological samples for each segment was performed using ImageJ on sequential 6 μm sections obtained at 300 μm intervals.

### Immunohistochemistry of murine atherosclerotic plaques

Frozen cryosections sections were thawed and washed twice in PBST (PBS with 0.01% Tween) before being blocked with endogenous 3% peroxidase (H_2_O_2_) for 20 min at room temperature. After two washing steps in PBST, the samples were blocked in 10% normal horse serum for 30 min at room temperature. Sections were washed three times and blocked with an Avidin/Biotin Blocking Kit (Vector Laboratories, Newark, CA, USA) for 30 min at room temperature before being rinsed in PBST and incubated with primary antibodies overnight at 4 °C. The primary antibodies used were anti-TER-119 (rat anti-mouse, cat 13-5921-82, 0.5 mg/mL, Thermo Fisher Scientific, Waltham, MA, USA). After antibody staining, sections were washed three times in PBST. Detection was achieved using a Vectastain ABC kit and DAB substrate (Vector Laboratories) and counterstaining with Mayer’s Haematoxylin for 15 s. Samples were dehydrated with an ethanol gradient, cleared in xylene and mounted with DEPEX. For visual clarity, the brightness and contrast were adjusted identically across the representative images shown in Figure 4 using Adobe Lightroom.

### Fluorescence-immunohistochemistry analysis of murine atherosclerosis

Frozen tissue sections were thawed, washed and permeabilised with 0.1% Triton X-100 for 10 min before being blocked with 1% BSA in PBST (0.05% Tween) for 45 min at room temperature. Following blocking, sections were incubated with fluorophore-conjugated antibodies for 1 h at room temperature. Antibodies used were Gr-1 (Ly6G/Ly6C)-AF488 (rat anti-mouse, cat 53-5931-82, 0.5 mg/mL, Thermo Fisher Scientific), anti-MHC II (I-A/I-E)-AF594 (rat anti-mouse, cat 107650, 0.2 mg/mL, BioLegend) and CD68-eFluor660 (rat anti-mouse, cat 50-0681-82, 0.5 mg/mL, Thermo Fisher Scientific). After antibody staining, sections were washed three times with PBST and stained with Hoechst 33342 (Abcam, Cambridge, UK) for 5 min, followed with two washes and a quick dip in distilled water before being mounted with Prolong Diamond Antifade (Invitrogen). Spleen tissue sections were used as positive controls to validate the staining specificity. Negative carotid controls were included and stained according to the above protocol, substituting primary antibodies with 1% BSA for the Gr-1 and anti-MHC II antibodies, and an IgG-eFluor660 isotype control (rat anti-mouse, cat 50-4321-82, 0.5 mg/mL, Thermo Fisher Scientific) for the CD68. Samples were imaged using a Zeiss LSM880 microscope and processed using the Zeiss Zen software. Images were analysed using ImageJ incorporating StarDist, a cell/nucleus detection method for microscopy images.^103^ To match the ImageJ analysis output colours to the immunofluorescence colourings available in Zen, the “replace colour” tool in Adobe Photoshop was applied for presentation in figures.

### AtheroExpress Biobank Study

The AtheroExpress Biobank Study (www.atheroexpress.nl) is an ongoing biobank that includes extensive baseline characteristics, blood samples and atherosclerotic plaque specimens along with three years of clinical follow-up, as described in detail elsewhere.^89^ Directly relevant to this work, we assessed the matched histological and RNA-sequencing data from the plaques of 1093 patients undergoing carotid endarterectomy between 2002 and 2017.^104^ **Supplementary Table 13** summarises patient characteristics. This was performed in line with the Declaration of Helsinki and informed consent was provided by all study participants after approval for this study by the medical ethical committees of the various hospitals (University Medical Center Utrecht, Utrecht, The Netherlands, and St Antonius Hospital, Nieuwegein, The Netherlands) was obtained and registered under number 22/088.

### AtheroExpress Biobank sample collection, data processing

Carotid plaque specimens were removed during surgery and immediately processed in the laboratory following standardised protocols. Specimens were cut transversely into segments of 5 mm. The culprit lesion (the region with the most severe stenosis) was identified, fixed in 4% formaldehyde, embedded in paraffin and processed for histological examination. The remaining segments were stored at −80 °. We described the standardised (immuno)histochemical analysis protocols used in the AE previously.^89^ In short, 10-micron cross-sections of the paraffin-embedded segments were cut using a microtome, stained using standardised protocols and examined under a bright-light microscope. IPH was scored as no/yes using an H&E staining; lipid content (Lipid) was defined as no or <10% vs. >10%; macrophage content and SMC content were defined as no or minor positive staining versus moderate to heavy positive staining (Mac and SMC, respectively).^105,106^ All histological observations were performed by the same dedicated technician and interobserver analyses have been reported previously.^107^

To obtain matched bulk RNA-seq data, atherosclerotic plaques were ground while frozen with liquid nitrogen and then TriPure (Roche, Cat# 11667165001) was added and the plaque pieces were further disrupted using a Precellys 25 homogeniser (Bertin Instruments).^90^ Samples were incubated at room temperature for 5 min and centrifuged at 20,000 g for 1 min at 4 °C. The supernatant was mixed with chloroform and incubated at room temperature for 15 min. The sample was centrifuged at 12,000 g for 5 min at 4 °C and the upper phase was used for RNA isolation. Then, isopropanol and GlycoBlue (Invitrogen, Cat# 10301575) were added to the aqueous phase to precipitate the RNA and the mixture centrifuged at 12,000 g for 10 min at 4 °C. The pellets were washed with 75% ethanol and resuspended in RNase-free water.

The RNA isolated from archived advanced atherosclerotic lesions is often fragmented. To overcome this, CEL-seq2 was employed.^108^ CEL-seq2 yields the highest mappability reads to the annotated genes compared to other library-preparation protocols. The methodology captures the 3′ end of polyadenylated RNA species and includes unique molecular identifiers (UMIs), allowing direct counting of unique RNA molecules in each sample. Then, 50 ng of total RNA was precipitated using isopropanol and washed with 75% ethanol. After removing ethanol and air-drying the pellets, a primer mix containing 5 ng of primer per reaction was added, initiating primer annealing at 65 °C for 5 min. Reverse transcription was then performed: first-strand reaction for 1 h at 42 °C; heat-inactivated for 10 min at 70 °C; second-strand reaction for 2 h at 16 °C; and then put on ice until proceeding to sample pooling. The primer used for this initial reverse-transcription reaction was designed as follows: an anchored polyT, a unique 6-bp barcode, a UMI of 6 bp, the 5′ Illumina adapter and a T7 promoter, as described. Each sample now contained its own unique barcode due to the primer used in the RNA amplification, making it possible to pool together complementary DNA (cDNA) samples at seven samples per pool. cDNA was cleaned using AMPure XP beads (Beckman Coulter, Cat# A63882), washed with 80% ethanol and resuspended in water before proceeding to the in vitro transcription (IVT) reaction (Thermo Fisher Scientific, Cat# AM1334) and incubation at 37 °C for 13 h. Next, primers were removed via treatment with Exo-SAP (Affymetrix, Thermo Fisher Scientific, Cat# 78201.1.ML) and amplified RNA (aRNA) was fragmented and then cleaned with RNAClean XP (Beckman-Coulter, Cat# A63987), washed with 70% ethanol, air dried and resuspended in water. After removing the beads using a magnetic stand, RNA yield and quality in the suspension were checked using Bioanalyzer (Agilent Technologies).

cDNA library construction was then initiated by performing a reverse-transcription reaction using SuperScript II reverse transcriptase (Invitrogen/Thermo Fisher Scientific, Cat# 18064022) according to the manufacturer’s protocol, adding randomhexRT primer as a random primer. Next, polymerase chain reaction (PCR) amplification was performed using Phusion High-Fidelity PCR Master Mix with HF buffer (New England Biolabs, F531L) and a unique indexed RNA PCR primer (Illumina) per reaction, for a total of 11–15 cycles, depending on aRNA concentration, with 30 s of elongation time. PCR products were cleaned twice with AMPure XP beads (Beckman Coulter, Cat# A63882). Library cDNA yield and quality were checked using Qubit fluorometric quantification (Thermo Fisher Scientific, Cat# Q32851) and Bioanalyzer (Agilent Technologies), respectively. This process occurred across two independent experiments: AtheroExpress RNA Study 1 and 2 (AERNAS1: n=622; AERNAS2: n=471). Libraries were sequenced on the Illumina NextSeq 500 platform, paired end, 2 x 75 bp.

### AtheroExpress analysis

All code used in these analyses will be provided upon publication on GitHub (details below). To calculate a gene-module score (GMS) using the bulk RNAseq data, a modified version of the Seurat function AddModuleScore was used. To test for the association of GMS in relation to the plaque instability index and individual plaque phenotypes (Lipid, Mac, IPH, SMC), we used ordinal and logistic regression as appropriate. We corrected for age, gender, year of surgery, hypertension status, diabetes status, current smoker status, lipid-lowering drugs (LLDs), antiplatelet medication, eGFR (MDRD), BMI, medical history of CVD (which is a combination of coronary artery disease history, stroke history and peripheral interventions) and stenosis. To investigate the association between the GMS and ischemic stroke during follow-up (time-to-event analysis), we performed survival analysis using Kaplan–Meier estimation and Cox proportional hazards models in R. Using the corresponding follow-up time associated with ischemic stroke, we stratified individuals based on the median value of the GMS. These strata (low vs. high expression) were used to generate Kaplan–Meier survival curves using the survfit() function and visualised with risk tables and stratified log-rank comparisons. Next, Cox proportional hazards models were fitted using the coxph() function to estimate hazard ratios (HRs) for each GMS stratum, adjusted as above.

### Statistical analysis (excluding scRNA-seq and AtheroExpress analysis)

All statistical analyses were conducted using GraphPad Prism (Version 10.2.3). Outlier testing was conducted using the ROUT test followed by normality testing using Shapiro–Wilk. Where there was normal distribution: for those with equal variance (defined by the F test), an unpaired Student’s t test was used; in the absence of equal variance, a Welch’s t test was used. In the absence of normality, a Mann–Whitney test was used. For data with more than two groups: normally distributed data with equal variance (defined by a Brown–Forsyth test) were assessed using a one-way ANOVA with Tukey’s multiple comparison test; normally distributed data with non-equal variance were assessed using a Brown–Forsythe and Welch ANOVA with Dunnett’s T3 multiple comparisons tests; non-normally distributed data were assessed using a Kruskal–Wallis with Dunn’s multiple comparisons test. For grouped data, a two-way ANOVA was used. For correlative analyses, data were binned into 50 intervals and then analysed using linear regression. For all, a p value of <0.05 was considered statistically significant.

## Data availability

The scRNA-seq data will be deposited on GEO and made publicly available upon publication. Raw RNA-seq data and (normalised) count matrixes with associated clinical and histological data from the AtheroExpress Biobank Study are not publicly available due to research participant privacy and consent. There are also restrictions on use by commercial parties and on open sharing based on (inter)national laws and regulations and written informed consent. Consequently, these data (and additional clinical data) are available only upon discussion and upon signing a data-sharing agreement via DataverseNL at this address: https://doi.org/10.34894/D1MDKL and within a specially designed environment provided by UMC Utrecht. However, part of the single-cell RNAseq from AtheroExpress is available through www.plaqview.com. The publicly available scRNA-seq data analysed in this study can be accessed through GEO (GSE20658).

## Code availability

The code used in this manuscript will be made available upon publication. Code used for scRNA-seq analysis and figure production can be found at https://github.com/JonathanNoonan/Noonan-CD8Athero. The SCENIC analysis can be found at https://github.com/pinto-lab/Noonan-CD8Tcell. The code used to analyse the data from the AtheroExpress Biobank Study is available at https://github.com/CirculatoryHealth/Noonan-CD8Athero.

## Acknowledgements

The authors thank AMREP Animal Services, the Baker Institute Flow Cytometry Core, the Alfred Research Alliance Animal Ethics Committee and the Alfred Research Alliance Flow Cytometry Core for their support of this research. The authors also thank the late Dr Nicholas Wong for his support with bioinformatics analyses and recognise his immense contributions to the research community. We are thankful for the support of the Leducq Foundation’s “PlaqOmics” and “AtheroGen”, and the Chan Zuckerberg Initiative’s “MetaPlaq”. The research for this contribution was made possible by the AI for Health working group of the EWUU alliance (https://aiforhealth.ewuu.nl/). The collaborative project Getting the Perfect Image was co-financed through Health∼Holland’s (Top Sector Life Sciences & Health) PPP Allowance. We would like to thank all those involved in the AtheroExpress Biobank Study, including the Departments of Surgery at the St Antonius Hospital Nieuwegein and University Medical Center Utrecht. We would like to thank all participants in the AtheroExpress Biobank and Alfred Hospital studies; without you, these kinds of studies would not be possible.

